# Development of a PEGylated Parylene Nanopocket Membrane for the Capture and Release of Lipid Vesicles

**DOI:** 10.1101/2025.06.02.657433

**Authors:** Munther Alsudais, Kevin E. Petersen, Atiyeh Hosseinifakhr, Leanna H. Frasch, Shayan Gholizadeh, Mehdi Dehghani, Thomas R. Gaborski

## Abstract

Membranes are commonly used for the separation and purification of a variety of biological species. In this study, we developed a nanopocket membrane that can capture nanoparticles and extracellular vesicles in tangential flow filtration by pulling the species of interest into a nanopocket, while tangential flow washes away particles too large to be captured. We developed the pores using a four-step lithography process. By optimizing photolithography parameters, particularly exposure dose, we achieved well-defined pore shapes with the desired wall tilt, forming nanopocket geometries. We validated membrane performance using bead mixtures of varying diameters (50 nm –10 µm), demonstrating its ability to pass small particles, capture mid-sized particles, and exclude larger ones. Surface modification with poly(l-lysine)- grafted poly(ethylene glycol) enhanced this performance by reducing pore size and improving particle release. Additionally, we demonstrated selective capture and release of liposomes as well as EVs in the presence of fluorescently labeled BSA (f-BSA) protein. During this process, EVs were retained and later released, while f-BSA passed through, confirming selective capture and reduced protein contamination. These PEGylated nanopocket membranes constitute a low-pressure platform for gentle, size-selective isolation of nanoscale bioparticles, laying the groundwork for improved extracellular-vesicle analytics and point-of-care diagnostic applications.

## Introduction

Extracellular vesicles (EVs) are cell-derived lipid nanovesicles that have attracted widespread interest for diagnostics and therapeutics, yet isolating EVs from biological fluids remains challenging. Conventional ultracentrifugation, the most common isolation method, is time-consuming and can co-isolate abundant non-vesicular macromolecules or even induce EV aggregation^1–3^. Membrane ultrafiltration offers a size-based alternative, but biological samples tend to foul the filters. Proteins and other contaminants readily form deposits or adsorb to the membrane, causing significant EV loss and reduced yield^2^. These issues motivate the development of more efficient, gentle EV separation techniques. One proven strategy to mitigate fouling is the use of tangential flow filtration (TFF), where fluid flows parallel to the membrane surface to continuously sweep away buildup. TFF-based systems provide continuous filtration with improved scalability and clog-prevention compared to normal (dead-end) filtration^4^, and have shown superior EV recovery and purity relative to ultracentrifugation in several studies.

While porous membranes have proven highly effective for general size-based separation, recent advancements have led to the development of more structurally refined designs. One such innovation is the nanopocket membrane, engineered to enhance the precision of particle capture and controlled release based on size. These membranes are characterized by a conical geometry, in which the pores gradually narrow from the top to the bottom, enabling size-selective retention and passage of target particles. This structural specificity makes nanopocket membranes particularly valuable in biomedical engineering. Researchers have focused on optimizing nanopocket membrane fabrication and characterizing their structural properties ^5–8^, while others have applied them in various contexts, including the capture-and-release of synthetic particles and biofluids such as proteins and cells ^9–12^. Despite the promising improvements offered by nanopocket membranes, a key limitation persists: they are typically fabricated with thicknesses exceeding 5 µm, leading to higher operating pressures and reduced separation efficiency for smaller particles due to the increased likelihood of trapping within long pore channels. ^13^. Recent studies have demonstrated that ultrathin membranes can operate effectively at significantly lower pressures compared to their thicker counterparts ^13,14^. Moreover, the reduced thickness shortens the pore channel, allowing particles to pass through more easily and with less risk of trapping. These findings highlight the potential benefits of integrating ultrathin architectures into nanopocket membrane designs. Consequently, leveraging ultrathin membrane fabrication techniques to develop next-generation nanopocket membranes with thicknesses below 2 µm offers a valuable strategy for improving performance under reduced pressure conditions.

In our previous work, we developed ultrathin silicon-based nanoporous membranes with straight-walled pores for various applications, including separations ^13,15–18^. We extended the utility of these ultrathin silicon nanomembranes by developing scaffolding and liftoff approaches^19^. To reduce fabrication cost and complexity and further increase membrane surface area, we researched fabrication of ultrathin porous parylene C (UPP) membranes for biological lab-on-a-chip applications ^20,21^. In this study we further refined these membranes by fabricating nanopockets using advanced lithographic techniques. To optimize membrane performance for biomedical applications, we modified the surface using poly(L-lysine)- poly(ethylene glycol) (PLL-PEG). Drawing on our prior work with PLL-PEG functionalization ^22^, this modification was designed to minimize non-specific binding and tune pore accessibility for target isolation. We chose PLL-PEG due to its well-documented protein-repellent properties via steric repulsion ^23^. We systematically evaluated membrane performance through dead-end and tangential flow experiments using an H-shaped microfluidic channel with HPLC fittings, enabling direct fluorescence microscopy visualization with in-line pressure measurements. The system successfully demonstrated capture and release of target particles including beads, liposomes, and extracellular vesicles.

## Results and Discussion

In previous works ^15,16,24,25^, ultrathin nanomembranes with straight pore walls were utilized (Figure 1a). However, this study introduces a membrane with a pore geometry design, referred to as nanopockets, where the pores gradually narrow to enhance size-based particle capture and release (Figure 1b). This design selectively captures particles with diameters between those of the larger and smaller pore openings as these particles enter and become trapped within the nanopockets. Meanwhile, tangential flow effectively washes away particles that are too large to enter the nanopockets while diameters smaller than the lower pore openings pass through the membrane without obstruction.

**Figure 1.**
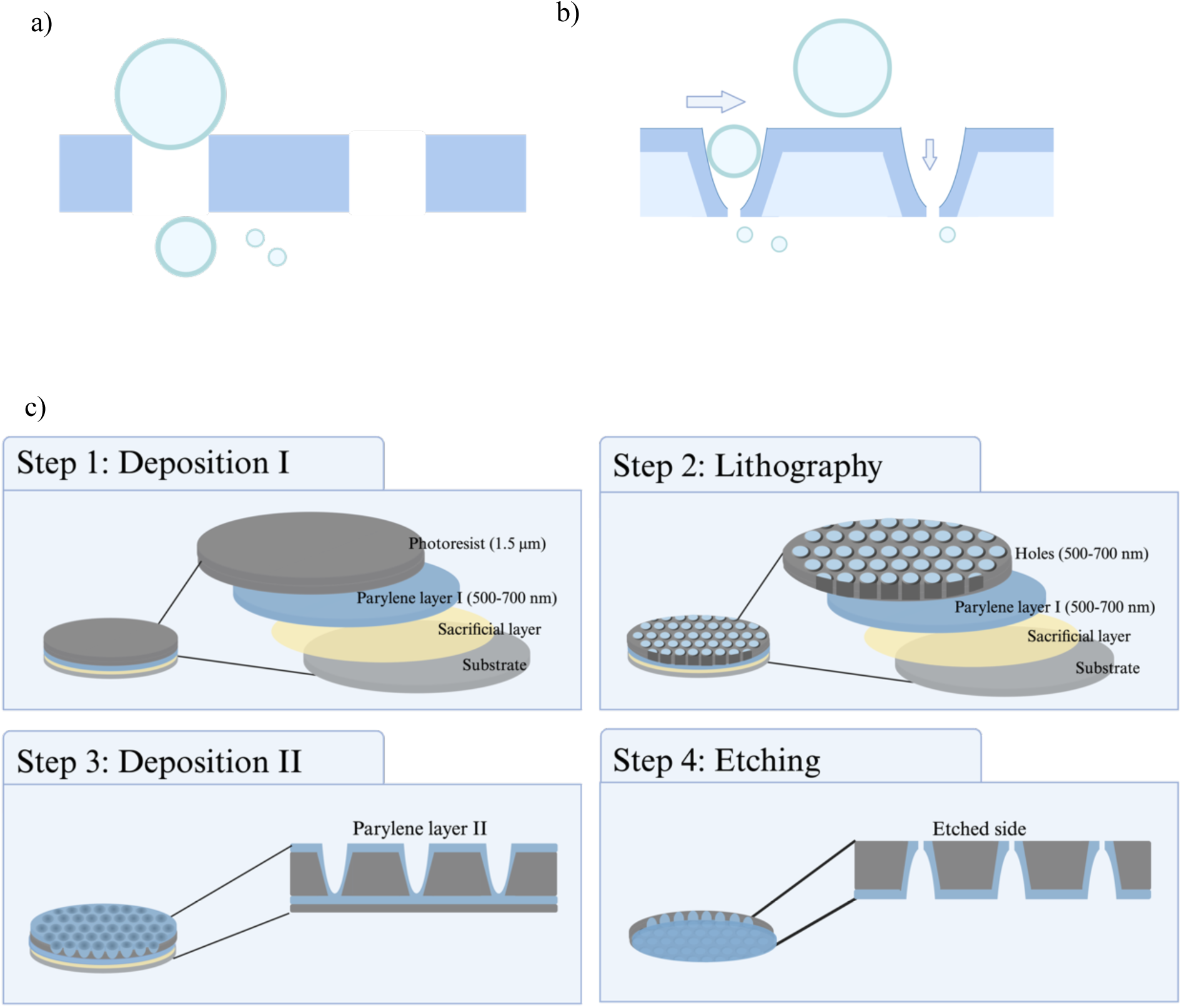
Overview of membrane configurations and fabrication steps. a) Previous studies utilized ultrathin nanomembranes with straight pore walls ^14,33,42,43^. b) Illustration of the ultimate design of the nanopocket membrane. This design features a top pore size larger than the bottom pore size, with the membrane thickness determined by the amount of parylene deposited on top (represented in dark blue) and the thickness of the photoresist (represented in light blue). c) Fabrication process for a nanopocket geometry membrane: **Step 1: Deposition I** - A water-soluble sacrificial layer, parylene layer I, and photoresist layer are deposited. **Step 2: Lithography** - Patterned holes with diameters ranging from 500 to 700 nm are created in the photoresist layer. **Step 3: Deposition II** - the deposition of parylene layer II results in the final nanopocket shape. **Step 4: Etching** - after the membrane is attached to the microfluidic device, the bottom was partially etched using reactive ion etching (RIE).

### Nanopocket membrane fabrication

The fabrication process for the nanopocket membrane is illustrated in Figure 1c, and consists of four main steps: deposition I, lithography, deposition II, and etching. The resulting nanopockets had a wide opening at the top, measuring between 500 and 700 nm, which tapers to a smaller opening at the bottom, ranging from 130 to 170 nm. The thickness of the membrane ranges from 1.5 to 2 µm, (Figure 1b).

#### Step 1: Deposition I

The deposition process involves applying three layers onto the 6-inch silicon (Si) wafer. Initially, a sacrificial layer was deposited to facilitate the subsequent removal of the membrane. This was followed by a parylene layer, referred to as parylene I in Figure 1c, which shields the sacrificial layer from the photoresist layer deposited over parylene I to pattern the holes.

To assess the uniformity of parylene deposition, three wafers were coated simultaneously in each run inside the parylene deposition chamber. The positions of the wafers on the chamber plates were fixed across all runs to minimize placement-related variation. To validate the deposited thickness, a NanoSpec meter (Nanometrics Model 210) configured with the known refractive index of parylene was used to measure the parylene thickness at the same location on all wafers (Figure S1). The parylene layer thickness ranged from 500 to 700 nm, with a standard deviation of less than 5% ^26^. This method ensured consistent and reliable results across multiple runs, as corroborated by other studies ^26^.

Reproducibility of the parylene layer thickness was assessed through wafer-to-wafer and run-to-run comparisons using a physically indexed template to map the measurement regions across the wafers. For wafer-to-wafer comparisons, three wafers from the same run were analyzed, with a total of 125 measurements taken per wafer. These measurements were distributed across a 5×5 grid of squares, with each square receiving five measurements (Figure S1). The 125 measurements from each of the three wafers were compared across two separate runs, involving a total of six wafers (Figures S2a and S2b). No significant differences were found between most wafers. In the run-to-run comparisons, we compared wafers from one run to those from another, analyzing the average measurements (n = 3 per run). As illustrated in Figure S2c, no significant differences were observed.

The photoresist layer was deposited using an SVG photolithography system to ensure uniform, unblocked pore formation. We attempted 1 and 1.3 µm thicknesses, but in order to minimize cracking and ultimate failure, we deposited 1.5 µm for all further membrane development. Thickness verification was performed using a NanoSpec meter, employing the known refractive index of the photoresist. Measurements taken across 15 independent wafers (n = 5 measurements per wafer) yielded an average thickness of 1.519 µm ± 0.038 µm, confirming consistent deposition.

#### Step 2: Lithography

We conducted experiments to determine the optimal exposure dose for achieving well-defined pore structures. To evaluate this, we used a previously developed mask with pore sizes of 0.5 µm ^21^. The focus offset was kept at zero while the exposure dose was varied from 380 mJ/cm² to 700 mJ/cm² in increments of 20 mJ/cm². These tests were conducted on a single wafer, which displayed a gradient of decreasing energy from one side to the other. Ultimately, we selected an exposure dose of 680 mJ/cm², as it effectively removed the photoresist and produced the desired tapering of the edges (Figure 2).

**Figure 2.**
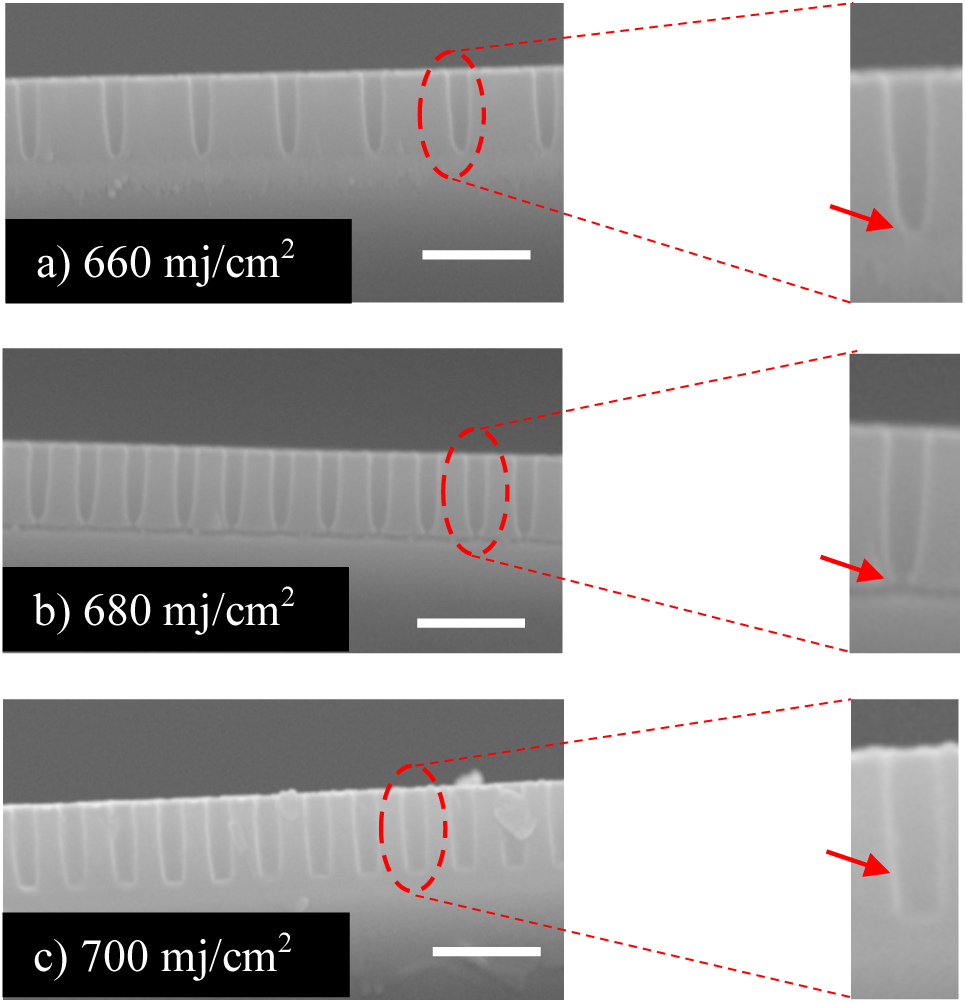
Various photoresist exposure doses and their effect on pore geometry: SEM images demonstrated exposure doses ranging from 660 mJ/cm² to 700 mJ/cm², incremented by 20 mJ/cm². Achieving tapered pore walls and complete removal of the photoresist at the bottom was crucial for obtaining the final nanopocket shape. a) Residual photoresist remained at the bottom of the pores as highlighted by the arrow. b) Tapered pore walls with complete photoresist removal at the bottom as indicated by the arrow. c) Straight pore walls indicated by the arrow. Scale bars = 2 µm.

#### Step 3: Deposition Parylene II

We investigated the effect of depositing parylene layer II on the pore size reduction and evaluated the consistency of deposition across different runs. Our focus was on the relationship between the amount of parylene used and the resulting pore diameter. The application of parylene layer II successfully narrowed the pore diameter, forming of the final nanopocket shape, which resembled a funnel.

To assess the uniformity of parylene deposition across various runs, we used different quantities of parylene (0.05, 0.1, 0.3, 0.5, and 0.7 g). The results (Figure S3) demonstrated a linear deposition process consistent with prior reports in the literature ^26–30^. These findings were crucial for evaluating the consistency and predictability of the deposition process, which are vital for ensuring the reliability and quality of the final product.

The impact of varying parylene amounts on nanopocket pore size was further studied using a single patterned wafer. With a patterned photoresist, different amounts of parylene (0.4 and 0.5 g) were applied to cause changes in the size of the nanopocket pores (Figure 3). It was noted that pore sizes decreased as the amount of parylene increased: without parylene the pore size was 915 +/- 25 nm, with 0.4 g it was reduced to 404 +/- 40 nm, and with 0.5 g it further decreased to 172 +/-42 nm.

**Figure 3:**
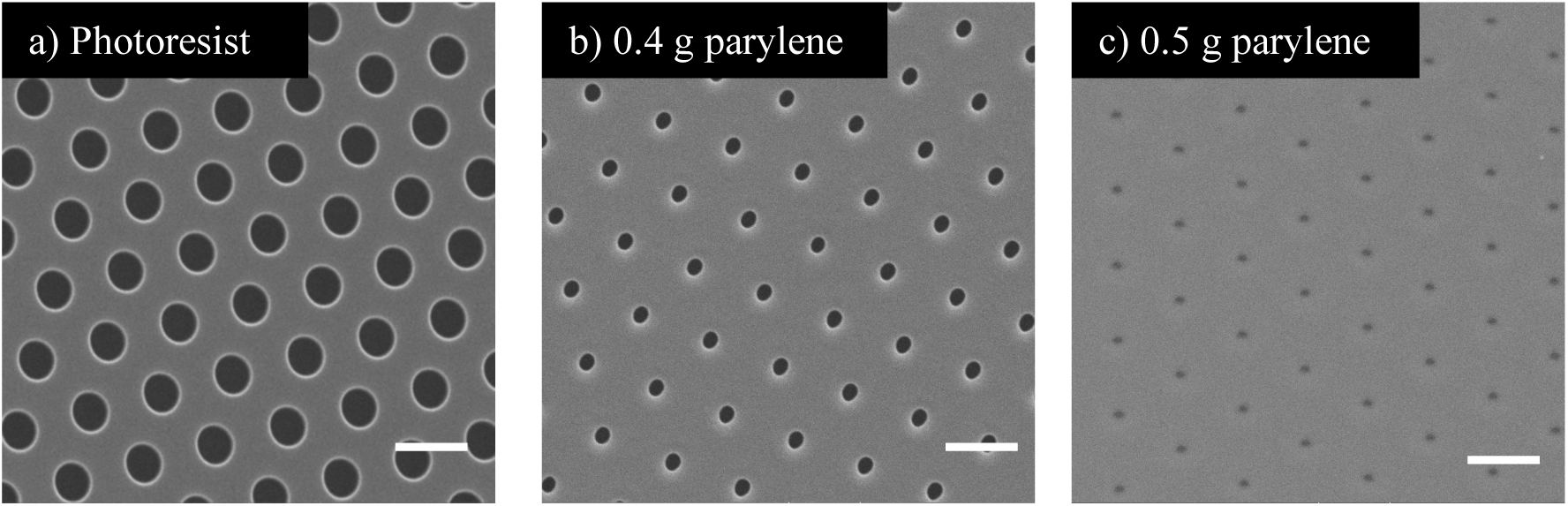
Effect of different parylene amounts on pore size. SEM images illustrated the reduction in pore size with increasing amounts of deposited parylene: a) Patterned photoresist layer without parylene, with an inner diameter (ID) of 915 (±25) nm, (n=3). b) 0.4 g of parylene applied on top the photoresist layer from (a), resulting in an ID of 404 (±40) nm, (n=3). c) 0.5 g of parylene on top of the photoresist layer from (a), achieving an ID of 172 (±43) nm, (n=3). Scale bars = 2µm.

#### Step 4: Etching

Prior to etching from the bottom, the wafer was sectioned into small pieces to facilitate integration onto an adhesive cover glass. The cover glass with the wafer piece was then immersed in deionized (DI) water. This step dissolved the sacrificial layer and released the membrane from the silicon chip while ensuring it remained attached to the cover glass.

A nanoforest ^31,32^ (wire-like) ^33^ phenomenon was encountered during the etching of parylene using oxygen (O_2_) (Figure S4). Previously, researchers have used various gas mixtures to address this issue. These mixtures included O_2_ ^34^ combined with argon (Ar) ^35^, O_2_ with fluoroform (CHF_3_) ^36^, O_2_ with sulfur hexafluoride (SF_6_) ^31^ and O_2_ with carbon tetrafluoride (CF_4_) ^33^.

To reduce the nanoforest, we examined different O_2_ gas flow rates (30 and 60 sccm). It was found that the nanoforest was almost the same with both flow rates (Figure S5 a and S5e). We then introduced the parylene along with low-flow O_2_, including Ar, CHF_3_, and SF_6_, at flow rates of 10 and 30 sccm. Among these gases, CHF_3_ and SF_6_ performed better at higher flow rates (figure S5). However, the etch rate was found to be unstable at high CHF_3_ and SF_6_ flow rates. It was determined that a stable flow rate of around 8 sccm was optimal for both CHF3 and SF6 gases.

To examine the variation in etching rates at different power levels (100 and 50 W), we conducted experiments using different samples. Each run included three devices placed in the etching machine chamber’s top, middle, and bottom positions. The etching times for the 100 W samples were 1, 2, 4, 5, and 7 minutes, while for the 50 W samples, the etching times were 2, 4, 5, 6, 8, 10, and 12 minutes. The data presented in Figure S6 demonstrates that etching at 50 W (109 nm/min) required more time compared to etching at 100 W (324 nm/min) to achieve the same thickness, highlighting the significant influence of power on the etching rate. Therefore, 100 W was selected as the power level for subsequent experiments.

To confirm the nanopocket geometry in pores, the membrane was inspected from different perspective as displayed in Figure 4. SEM revealed that the bottom pores were 130 - 170 nm in diameter, and the top-side pores were 500 - 700 nm in diameter. This confirmed the nanopocket shape.

**Figure 4:**
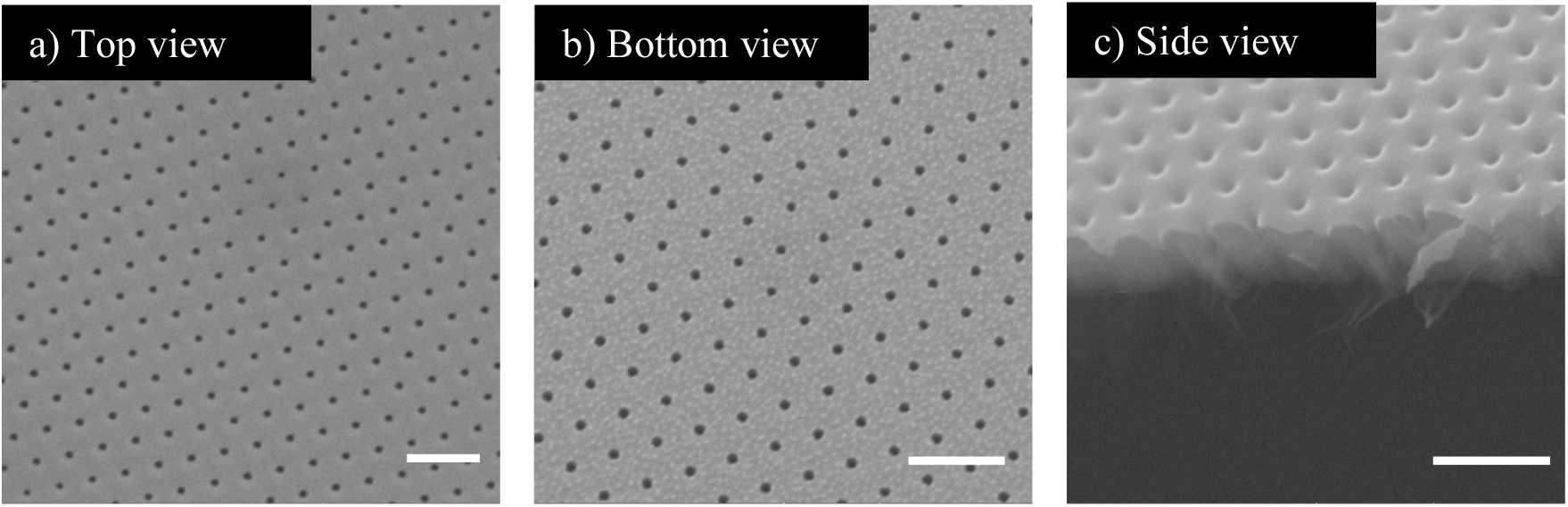
SEM images showing different perspectives of the membrane. Pore diameters differ on the two sides of the membrane, confirming successful fabrication of the nanopocket geometry. a) Top view of the membrane. b) Bottom view of the etched membrane, with pore sizes ranging from 130 to 170 nm. c) Side view highlighting the soft edge of the membrane and the top-side pores with diameters of ∼500 nm. Scale bars = 2 µm.

#### Step 5: PEGylation of Membranes

To modify the membrane surface to both improve non-fouling properties and further refine nanoparticle geometries, PEGylation was performed after device assembly by flowing the PLL-PEG solution through the system to ensure controlled grafting. Initial grafting conditions (PLL-PEG concentration, incubation time) were based on our previous work ^22^, with additional optimization experiments conducted to minimize protein fouling on parylene surfaces. Through systematic testing, we determined that a PLL-PEG concentration of 0.025 mg/mL successfully reduced protein adhesion (Figure S7).

To verify that PLL-PEG modification would not clog nanopockets, we analyzed polymer brush conformation. Although PLL-PEG’s chemical structure is well-characterized, predicting its spatial configuration remains challenging due to multiple interacting factors: copolymer weight ratio, grafting density, incubation time, and interfacial interactions (brush-substrate, brush-brush, and brush-solvent). We employed the Alexander-de Gennes ("blob") model as a theoretical framework for estimating brush length. This model predicts that when polymer chains are grafted above a critical density indicating chain overlap, they transition from a relaxed "mushroom" conformation to an extended brush-like state, stretching away from the surface ^37^.

Given the challenges in theoretical prediction of PLL-PEG brush length, we experimentally characterized brush conformation under our optimized grafting conditions (0.025 mg/mL) using atomic force microscopy (AFM). To avoid complications from membrane nanoporosity and wrinkling artifacts, we performed AFM measurements on flat parylene substrates under both dehydrated and hydrated conditions (Table S1). Our measurements revealed consistent brush lengths of approximately 33.16 ± 5.18 nm in the dehydrated state and 31.57 ± 4.52 nm in the hydrated state, demonstrating minimal conformational change (<5%) between conditions. This near-identical behavior suggests that the PLL-PEG brush maintains a relatively stable structure regardless of hydration state.

We measured the pressure differential (ΔP) across membranes using upstream and downstream sensors. Unmodified membranes showed ΔP = 0.9 psi at a 2 µl/min flow rate, while PEGylated membranes required ΔP = 2.5 psi at 0.5 µl/min (Figure 5a). Triplicate measurements showed consistent results. These values fall between ultrathin silicon nitride membranes (ΔP ≈ 0.2–0.3 psi) and thick track-etched membranes (ΔP ≈ 10 psi) ^13^.

**Figure 5.**
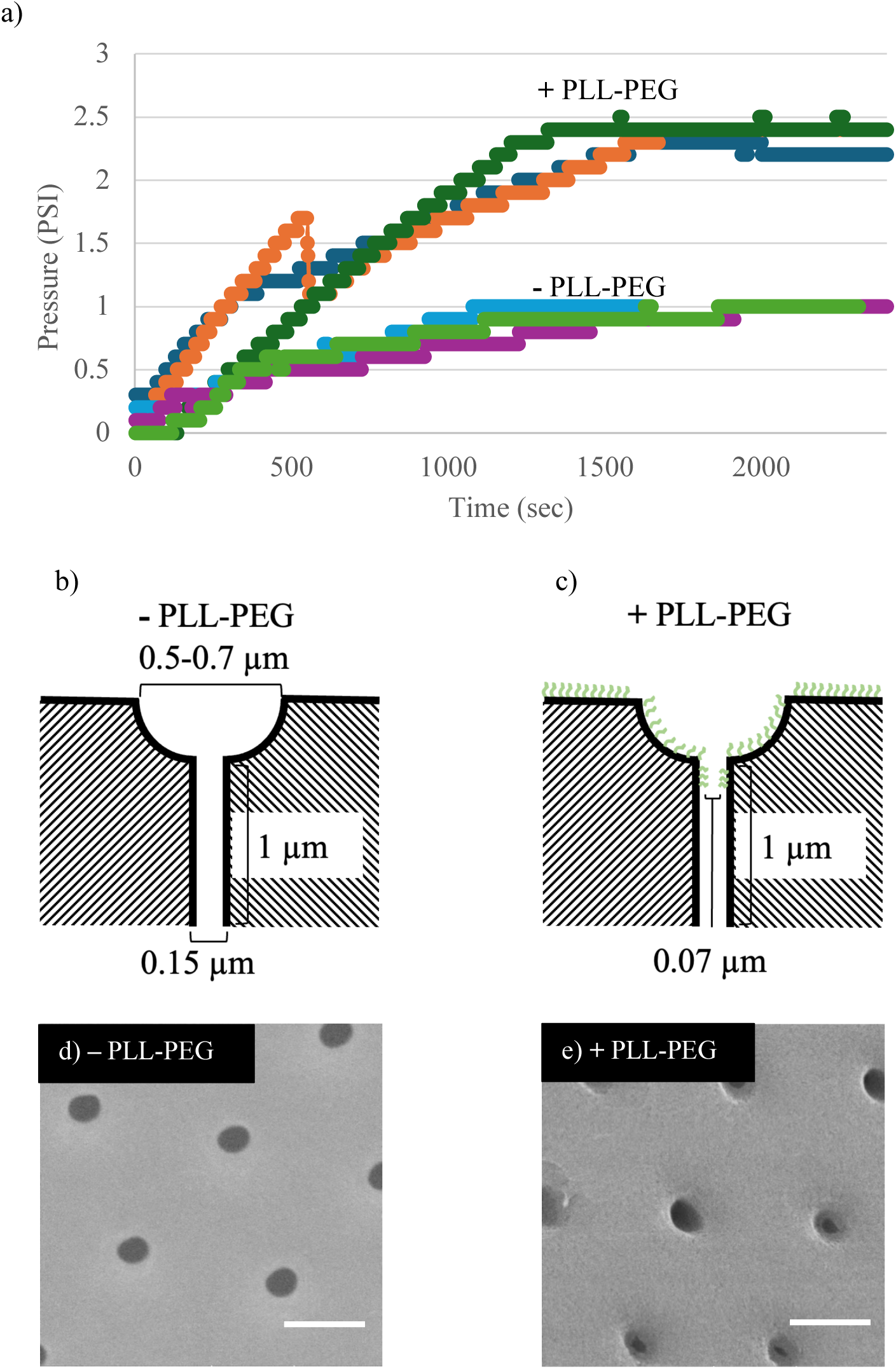
Characterization of PLL-PEG coating on parylene substrates. a) Permeability experiments showing ΔP measurements for three membranes with identical pore geometry. ΔP values stabilized at ∼1 psi for non-PEGylated membranes and ∼2.5 psi for PEGylated membranes. b) Nanopocket shape of a non-PEGylated membrane; membrane thickness was analyzed using Dagan’s equation, as shown. c) Hypothesized nanopocket shape of PEGylated membrane, demonstrating a reduced pore diameter. The PLL-PEG brushes are shown in green. d and e) SEM images comparing parylene membranes with and without PLL-PEG; scale bars = 500 µm.

Using the Dagan equation (Eq. 1) with a pore density of 8.5 × 10⁵ and a pore diameter of 130 nm (Figure 4b), we calculated ∼1 µm thickness of membrane region below the nanopocket, consistent with experimental measurements (Figure 4c). This agreement supported using the model for pore modification studies. Applying the Dagan equation to PEGylated membranes (ΔP = 2.5 psi at 0.5 µl/min, thickness = 1 µm) yielded a reduced effective pore diameter of ∼70 nm (Figure 5c). The approximately 60 nm diameter decrease aligns with AFM-measured PLL-PEG brush lengths of 31.6-33.2 nm, suggesting uniform deposition on both flat and porous surfaces. SEM images confirmed this through characteristic ring structures in modified membranes (Figures 5d and 5e).

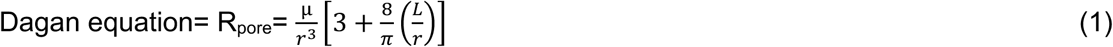

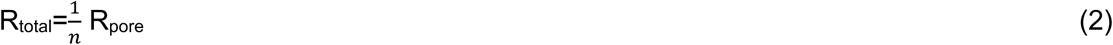

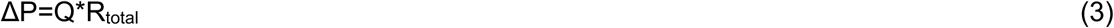

Viscosity “µ” [pa s^-1^], Pore radius “r” [m], Pore thickness “L” [m], Volumetric flow rate “Q”, Pressure “ΔP” [Pa], and Number of pores (n).

#### Assessment of Membrane Functionality

Our research group previously developed the Tangential Flow for Analyte Capture (TFAC) microfluidic device using a layer-by-layer assembly method utilizing flat nanoporous silicon nitride membranes ^13^. Our current design differs from previous work by positioning the sample inlet at the bottom and the filter channel at the top, with a parylene nanopocket membrane placed in between (Figure 6a).

**Figure 6:**
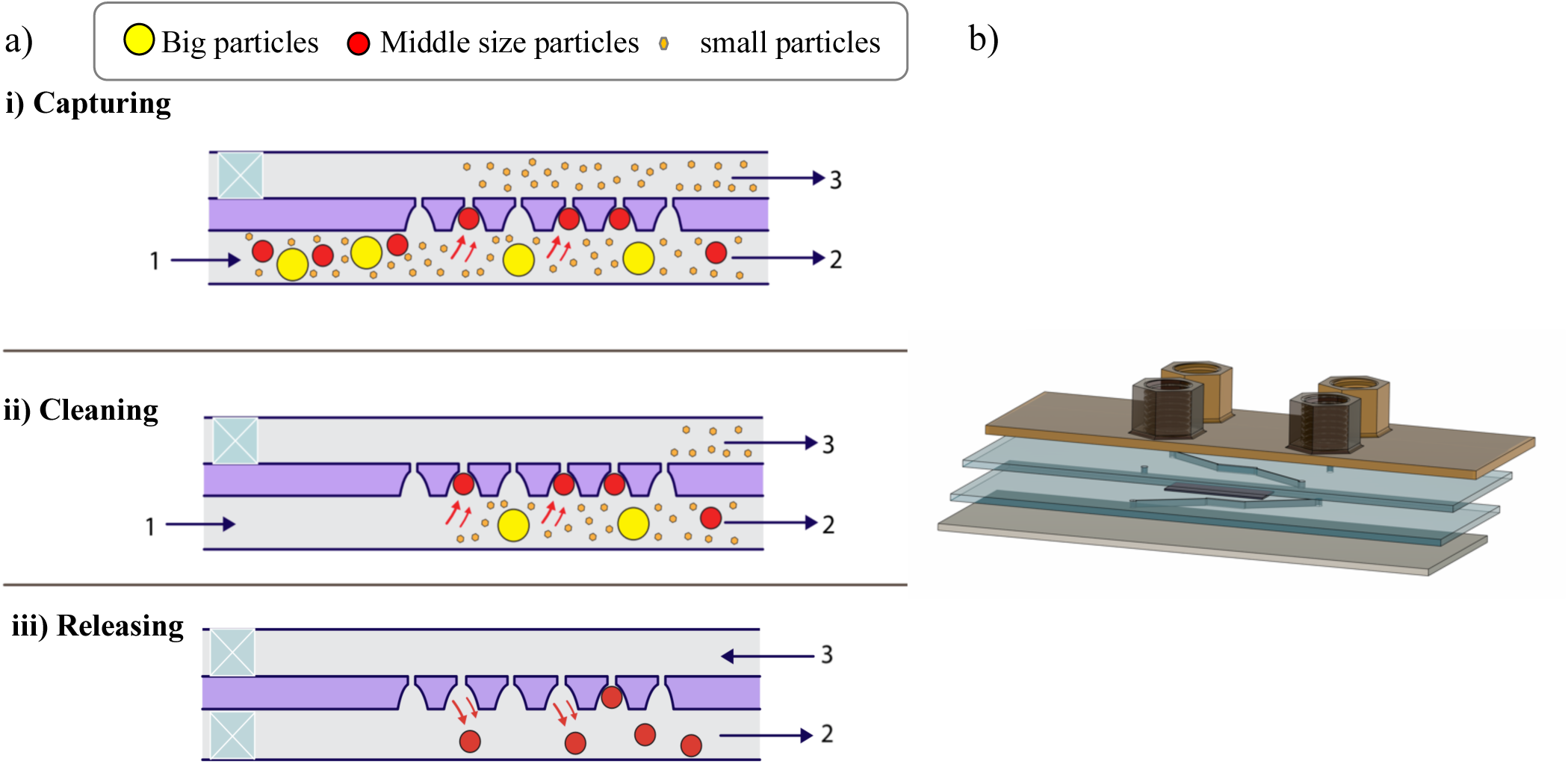
TFAC technique for particles capture and release. a) Assessing nanopocket membrane capture and release capabilities using fluorescent polystyrene beads/liposomes/EVs in the presence of proteins. i) Capturing Step: The sample is introduced through inlet 1 using a syringe pump towards outlets 2 and 3. A withdrawing syringe pump at outlet 3 drives particle motion toward the membrane, while outlet 2 remains open to air. ii) Cleaning Step: A clean buffer is then passed through inlet 1 under the same flow conditions as the capturing step to wash away all particles not trapped in the membrane. iii) Releasing Step: Captured particles are released by blocking inlet 1 and applying a reverse flow of 2 µL/min through the membrane. b) Microfluidic Device Design. The microfluidic device consists of four components: a 3D-printed layer, a nanopocket membrane sandwiched between two adhesive layers, and a cover glass.

The device consists of four layers: a 3D-printed layer, two adhesive layers, and a cover glass, all of which had matching dimensions (25 × 75 mm). The 3D-printed layer features four ports designed to be compatible with standard microfluidic tubing connectors. The two adhesive layers sandwich the membrane, with the larger pore opening size facing the cover glass. This arrangement enables gravity and sedimentation to reduce capture of large micron-scale particles such as cells and allowed for easy visualization of particle behavior during release using an inverted microscope.

To summarize the device assembly process, we followed this sequence of steps: First, we attached adhesive layer 1 to the 3D-printed layer. Then, we applied adhesive layer 2 to the cover glass. Next, we adhered a sectioned wafer featuring the nanopocket pattern to the cover glass, with the larger opening facing downward, and immersed it in DI water. This process dissolved the sacrificial layer, allowing the nearly complete membrane to detach from the silicon chip while remaining attached to the adhesive-covered glass. After etching the bottom of the membrane, we bonded the adhesive-covered 3D layer to the adhesive-covered glass with the membrane sandwiched in between (figure 6b).

To evaluate the functionality of the nanopocket membrane as a filter, we used the TFAC device to capture and release beads in three steps (Figure 7a): Capturing, Cleaning, and Releasing. For the capturing step, positive pressure was applied at the bottom channel inlet while negative pressure was applied at the top channel outlet using syringe pumps to drive the particle mixture towards the membrane. Smaller particles (50 nm beads used as a tracer and mimic of larger proteins or protein aggregates that could clog nanopores) either passed through the pores or exited the channel, while larger particles (10 µm particles to mimic cells) sedimented to the bottom of the channel. The targeted mid-size particles (500 nm particles used mimic liposomes and large EVs) were captured on the nanopocket membrane. These sizes were also chosen so they could be readily distinguished from one another in situ using a light microscope. During the cleaning step, ultrapure water was introduced through the bottom channel inlet under the same flow conditions as the capturing step. This process helped to wash the channel and membrane surfaces of any uncaptured beads. For the releasing step, the bottom channel inlet was closed, and positive pressure was then applied at the top channel outlet to release the targeted particles from the membrane, allowing them to be collected at the bottom channel outlet.

**Figure 7:**
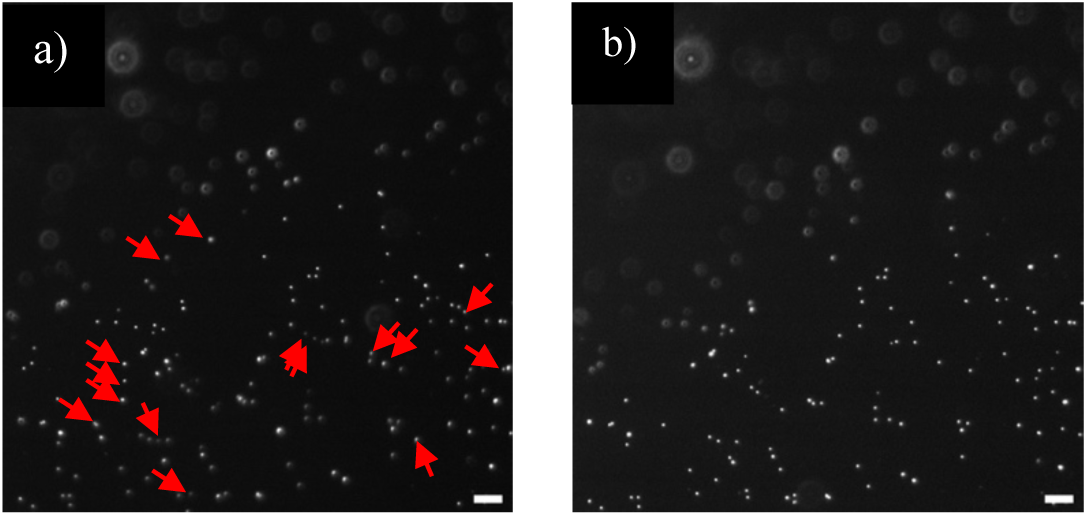
Isolation experiments using 0.05, 0.5, and 10 µm beads mixture. a and b) A 1:1:1 mixture of 0.05, 0.5, and 10 µm beads processed using an uncoated membrane, a) the membrane captured 0.5 µm beads after the capturing step, the arrows pointed on the beads been released, b) the membrane released some 0.5 µm beads after the releasing step. Scale bars = 10 µm.

#### Non-PEGylated Nanopocket Membrane: Bead Separations

To confirm isolation based on particle size, beads of varying diameters (50 nm, 500 nm, and 10 µm) were tested. The 50 nm beads successfully passed through the membrane and were detected in the filtrate collected from outlet 3, as confirmed by fluorescence imaging on a glass slide (Figure S8 a and S8d). This observation was crucial for understanding the membrane’s selectivity in allowing smaller particles to pass through. In contrast, 500 nm beads, were captured by the membrane, confirming its ability to retain particles larger than the reduced pore threshold (Figures S9 a and S9b). Notably, the 10 µm beads were not present on the membrane surface, likely due to sedimentation occurring before they could contact the membrane (Figures S9 c and S9d). This visually confirmed the membrane’s selective filtration capabilities and underscored its potential for nanoparticle and EV isolation. The differences in these outcomes emphasized the membrane’s dual functionality: it allows the passage of smaller particles while efficiently capturing targeted beads and preventing the adherence of larger beads.

To further investigate the membrane’s ability to isolate and release mixed populations of particles, we tested a 1:1:1 mixture of beads with sizes of 50 nm, 500 nm, and 10 µm. After introducing this mixture, we found that the 0.5 µm beads were selectively captured and released (Figure 7a and 7b). Meanwhile, the 10 µm beads were not detected on the surface, and the 0.05 µm beads either passed through the membrane or washed away down the bottom channel. This demonstrates the membrane’s effectiveness in size-based discrimination. Bead capture and spatial distribution were analyzed using fluorescence microscopy and SEM (Figure S10). Fluorescence imaging showed 500 nm beads arranged in a geometric pattern matching pore locations, which SEM confirmed as specific pore capture.

#### PEGylated Nanopocket Membrane: Bead Release and Separations

We next investigated the PEGylated nanopocket membrane. Based on fluorescent image analysis of the membrane in situ, the coated membrane showed a significantly improved release rate for the 0.5 µm beads, increasing from approximately 14% in the uncoated condition to about 45% in the coated condition. These findings confirm that the membrane can not only capture but also release targeted particles. Moreover, they demonstrate that surface modification enhances functional performance by facilitating the release of particles from the nanopocket structures.

We found that both 50 and 100 nm beads could pass through the pore at the bottom of the nanopocket in unPEGylated membranes. With PLL-PEG-coated membranes, we found that 50 nm beads passed through and were detected in the filtrate from outlet 3, as confirmed by fluorescence imaging on a glass slide (Figure S8b). In contrast, the 100 nm beads (similar in size to small EVs) were retained by the PEGylated membrane, as also verified by fluorescence imaging (Figure S8e). These results suggest that the PLL-PEG coating reduces the membrane’s effective pore size to below 100 nm and is consistent with our permeability measurements (Figure 5).

#### Liposomes Separation

While the bead experiments demonstrated successful capture and release of nanoparticles, we also evaluated the membrane’s performance with more biologically relevant samples. Liposomes were selected because their flexible lipid bilayer structure closely resembles that of EVs, making them a better model for studying interactions between the membrane and nanoparticles. Specifically, we found that 170 nm (Figure S11a) liposomes could be successfully captured and released from the PEGylated membrane (Figure 8a and 8b). It’s important to note that since the membrane does not remain completely flat during operation, some regions were out of focus. To address this, we restricted image analysis to a 5000 µm region where the liposomes and EVs were in focus and assumed uniform capture and release across the membrane surface.

**Figure 8:**
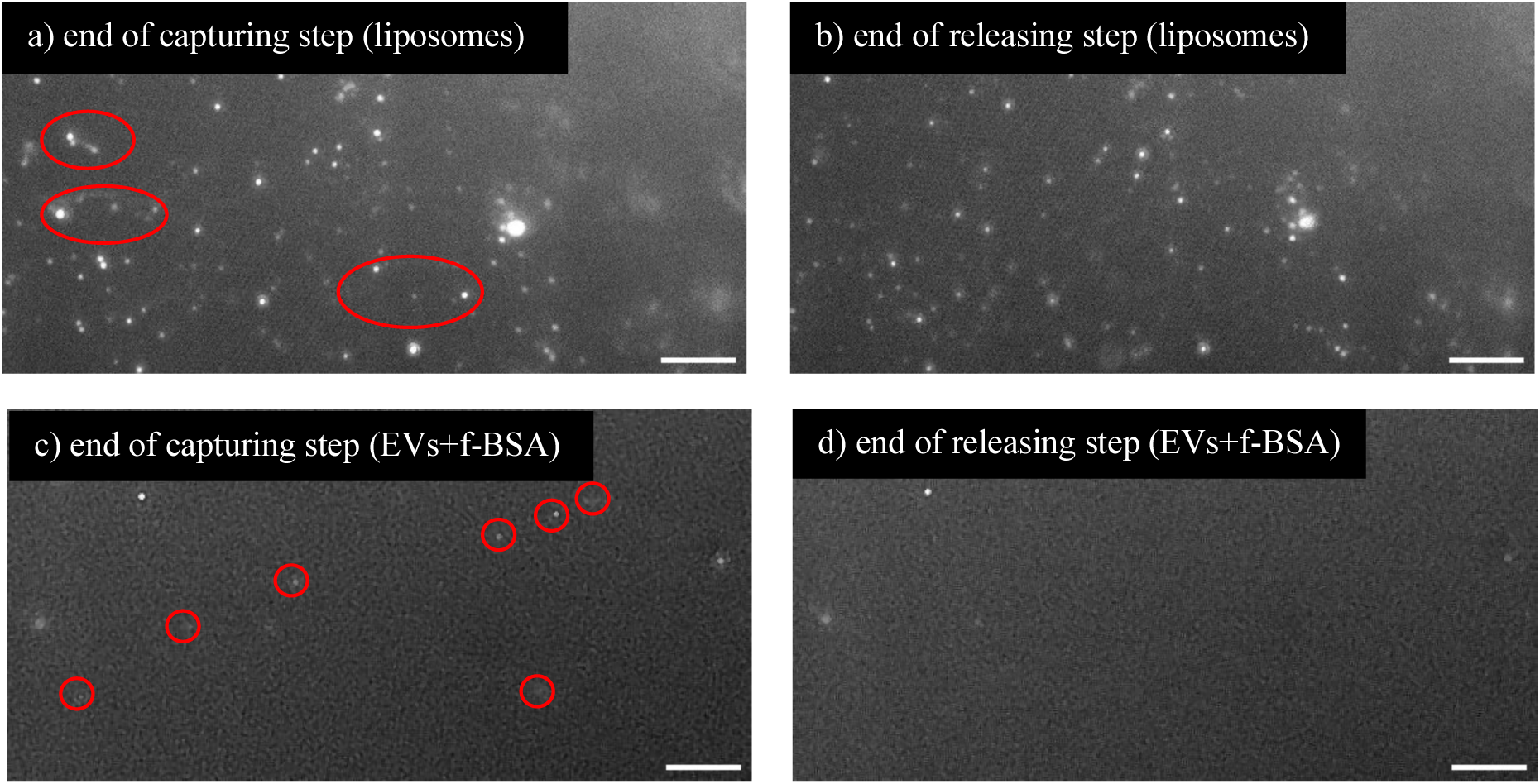
Isolation of liposomes and biofluid. a and b) Liposomes labeled with cell mask orange (CMO) were captured and released on a PLL-PEG–coated membrane, a) at the end of the capturing step, ovals indicated released liposomes, b) at the end of the releasing step. c and d) EVs in the presence of f-BSA were captured and released on a PLL-PEG–coated membrane, c) at the end of the capturing step, circles indicated released EVs, d) at the end of the releasing step. Scale bars= 10 µm

#### EV and f-BSA Separation

To evaluate the membrane’s performance with biologically relevant samples, we first tested human umbilical vein endothelial cell EVs, which have a more complex structure than liposomes. EVs contain a variety of transmembrane and surface bound proteins that affect their interaction with surfaces. These EVs were previously purified using conventional ultracentrifugation methods^1^. EVs (mean diameter: 145 nm, Figure S11b) were introduced into a device equipped with a PLL-PEG-modified membrane. The membrane successfully captured and released the EVs, confirming its potential for EV isolation (Figure 8 c and 8d). To determine whether smaller proteins could permeate the membrane, we introduced f-BSA and monitored fluorescence at outlet channel 3. The detection of f-BSA confirmed that the membrane’s pore size allowed selective transmission of proteins while retaining EVs (Figure S12a).

Next, we assessed the membrane’s ability to isolate EVs from a mixed sample containing both EVs and f-BSA (Figure 8c and 8d). During the capturing step, EVs were retained on the membrane surface, while f-BSA either passed through or was washed away (Figure 8c), demonstrating size-based exclusion of proteins. In releasing step experiments, EVs were liberated only after f-BSA clearance, ensuring contamination-free isolation (Figure 8d). Using in situ image analysis, we found the PEGylated membranes reliably released over 42% (±7.2) of captured EVs across three independent replicates, highlighting its effectiveness for selective particle recovery. These results underscore the membrane’s suitability for biofluid processing applications requiring high-purity, size-based separation. During imaging, we observed variability in the optical clarity of EVs on the membrane, with some appearing dimmer than others within the same frame. In contrast, liposomes prepared under similar conditions exhibited more uniform brightness. To confirm this observation, we immobilized EVs between a glass slide and a coverslip and found that the brightness differences persisted (Figure S12b). Fluorescent images of the membrane before and after these experiments suggested minimal if any f-BSA bound to the PEGylated parylene membrane. This is consistent with a conformal non-fouling PEG brush at the membrane surface.

## Conclusion

This study demonstrates the successful development of ultrathin (1.5 µm) parylene nanopocket membranes with tapered pores for size-selective biomedical nanoparticle separation and isolation. Optimized lithographic fabrication yielded reproducible pore geometries and tilted sidewalls, enabling low-pressure operation (∼1 psi). Post-assembly PLL-PEG grafting helped tune nanopocket and nanopore sizes. The membrane’s versatility was tested using polystyrene beads, liposomes, and EVs, which were all successfully captured and released even in protein-rich solutions. By combining ultrathin architecture, precise pore geometry, and antifouling surface modification, these membranes address critical biomedical challenges: gentle processing of sensitive biologics, size-selective filtration, and clogging resistance. Future work will focus on further refining nanopocket geometries. The current device may cradle EV-sized nanoparticles too strongly, making it difficult to release even when there is minimal interaction between the PEG coating and the nanoparticle. Additionally, we envision developing a microfluidic device with a series of nanopocket regions with increasing pocket size to capture and isolate different sized EV subpopulations.

## Methods

### Sacrificial Layer Deposition

We followed the protocol as previously published ^21^. A 4% v/v% solution of Micro-90 was used as a water-soluble sacrificial layer. This was achieved by spin-coating the solution onto a 6-inch (100) silicon wafer (500 rpm for 3 s, followed by 3000 rpm for 45 s). After all other layers were added to the wafer, the final step in the fabrication process was to expose the attached membrane to deionized (DI) water, which dissolved the Micro-90 and released the membrane from the wafer substrate.

### Parylene Coating

The parylene coating step has been previously published ^20,21^. Parylene-C coating was deposited using DPX-C dimer (Specialty Coating Systems, USA) in an SCS Labcoter 2 Parylene deposition system (PDS 2010, Specialty Coating Systems, USA). Silicon wafers were carefully positioned in the deposition chamber to ensure thickness uniformity across all runs. The dimer was loaded into the vaporizer. The process was initiated at a base chamber pressure of 10 mTorr, and the dimer-cracking furnace was heated to 690 °C. Subsequently, the vaporizer was ramped to a final temperature of 175 °C, leading to the sublimation of the dimer. The temperature ramp rate of the vaporizer was adjusted to maintain a chamber pressure of 25 mTorr. Thickness measurements were conducted using a NanoSpec Spectrophotometer (Nanometrics Incorporated, USA).

### I-Line Photolithography

We used the same protocol that had been published previously ^21^. All development, rinsing, and baking steps were preformed on the Silicon Valley Group (SVG) 88 Series Track. Hexamethyldisilazane (HMDS) prime from (MicroChemicals, Germany), served as an adhesion promoter between parylene and photoresist. The application involved spin-coating at 3000 rpm for 45 s, followed by baking at 140 °C for 1 min. AZ MiR 701 positive photoresist (MicroChemicals, Germany) was then applied through spin-coating at 1500 rpm for 45 s, followed by soft-baking at 90 °C for 1 min to achieve a 1.5 µm thickness. The spin-coating process followed the manufacturer’s recommendation of 500 rpm for 3 s, followed by 2200 rpm for 30 s.

The photoresist exposure was carried out using an ASML PAS 5000 i-line 5X Stepper (ASML, Netherlands) with final 0.5 µm pore sizes mask. The wafers underwent a post-exposure bake at 110 °C for 1 min. Subsequently, the exposed photoresist was developed using Microposit MF CD-26 developer (MicroChemicals, Germany), followed by hard-baking at 140 °C for 1 min.

### Reactive Ion Etching (RIE)

Reactive Ion Etching (RIE) was performed using the RIE-2000 (South Bay Technology Inc.) system. Several etching recipes were thoroughly evaluated to identify the most effective combination. The optimal gasses for this process were determined to be oxygen (O_2_) and Fluoroform (CHF_3_) (figure S5). The gas flows were precisely controlled at 30 standard cubic centimeters per minute (sccm) for O_2_ and 8 sccm for CHF_3_.

### Scanning Electron Microscope (SEM)

Samples were mounted and sputter coated with ∼3–7 nm of gold. Scanning electron micrographs were taken at different magnifications and an accelerating voltage of 20 kV using S-4000 SEM (Hitachi, Tokyo, Japan).

### Microfluidic device design and fabrication

The microfluidic device consisted of four layers: a 3D-printed layer, two layers of 3M adhesive, and a cover glass (Figure 5b). The 3D-printed layer, fabricated using Form 3B printer (Formlabs, USA) measured 75 × 25 mm, to be compatible with both the microscope platform and the cover glass. This layer features four ports designed to accommodate standard microfluidic tubing connectors, each with having a depth of 1 cm and a diameter of 0.8 cm. The ports were threaded using a 1/4NF28 tap (Pittsburgh) to ensure secure attachment of the connectors. Adhesive Layer 1 was applied to the 3D-printed layer to facilitate assembly.

The adhesive layers were cut from 468 MP adhesive (3M, purchased from Amazon part number: B074L5LSSZ, 2.3 mil (132 µm) thick with a 58 polycoated Kraft liner). The adhesive was laser cut with the plain side (non-3M logo side) up at 50% power, 100% speed, and one pass. The adhesive layers featured outer rectangular dimensions of 0.5 × 6 mm, internal circles 1.5 mm in diameter, four 10 mm-long channels.

A rectangular cover glass measuring 75 x 25 mm (Chemglass Life Sciences) was used to be compatible with the microscope’s working distance. Adhesive layer 2 was attached to the cover glass and folded along the dotted line with the membrane sandwiched between the adhesive layers 1 and 2.

### Bead Isolation Experiments

10 μm polystyrene green fluorescent beads (Thermo Scientific, USA), and 0.05 µm and 0.5 µm polystyrene red fluorescent beads (Magsphere, Canada) were used on the nanopocket membrane experiments. Flow experiments were conducted using two Chemyx Fusion 200 syringe pumps (Chemyx Inc., Stafford, TX). During the capturing step, a sample supply flow rate of 2 µl/min was applied, coupled with a pull rate of 1 µl/min. Subsequently, captured particles were released using a reversed flow of 2 µl/min through the membrane.

### PLL-PEG Grafting on Membranes

The microfluidic device was initially primed with ultra-pure water (2 µL/min flow rate, NE300 syringe pump) from port 1 to port 2. We then introduced an air bubble followed by injection of 0.1 mL of PLL-PEG solution (0.025 mg/mL in HEPES buffer; PLL(100)-g(28)-MPEG(44), Nanosoft Biotechnology) using a 7725 injection valve (Rheodyne LLC) at 2 µL/min until solution emergence at port 2. The system was then static-incubated for 30 minutes to allow polymer grafting, followed by thorough rinsing with ultra-pure water.

### Atomic Force Microscopy (AFM)

To assess the brush length of the PLL-PEG concentration and incubation time, the samples was assessed using two AFM systems, one for measuring brush length under dehydrated conditions and the other under hydrated conditions. A parylene coating was applied directly onto a silicon wafer as described above, followed by a 1.5 um layer of negative photoresist. (Mask 4 um pore size, 5% porosity).

The wafers were cut into 5×5mm samples, submerged in PLL-PEG solution and incubated for 30 minutes. Following incubation, the substrates were rinsed in deionized water, then dipped in acetone for 15 seconds to dissolve the photoresist and expose the PLL-PEG islands, followed by another rinse.

For brush length assessment under dehydrated conditions, a MultiMode 8-HR AFM (Bruker) with an SNL-10A cantilever was used under tapping mode. To assess the brush length under hydrated conditions, the samples were mounted onto a fluid cell chamber to which PBS was added. Surface scans were obtained using an MFP-3D AFM (Oxford Instruments) with TR800PSA cantilevers (Oxford Instruments) with a spring constant of 0.15 N/m in tapping mode.

For both methods, measurements were taken from three PLL-PEG islands on each sample and measured in duplicate to obtain height averages and standard deviations. Height measurements were calculated using the height profile outputs from measuring across the PLL-PEG islands.

### PLL-PEG Fouling Assessment

To assess suitable PLL-PEG concentrations, the anti-fouling properties with the concentration of grafting was of interest. Glass slides were cleaned with isopropyl alcohol and coated with parylene using the previously described recipe and system. Silicone gaskets were adhered to the slides and PLL-PEG at concentrations ranging from 0.025 to 0.9 mg/mL were added and incubated for 30 minutes, followed by a rinse with DI water. Then 0.1 mg/ml fluorescent bovine serum albumin, f-BSA (Invitrogen, OR, USA, Ref#A13100, Lot#2551334), was added to the gaskets and incubated for one hour at room temperature, followed by another rinse with DI water. The gaskets were filled with PBS and a coverslip was added on top. Using a fluorescence microscope (Leica DMI6000B), the slide surface of each gasket was imaged after which the relative brightness intensities were calculated and compared.

### Permeability experiments

The permeability experiments were conducted using the PressureMAT model PMAT4R system by (PendoTECH, USA). In the microfluidic device, two pressure sensors, P_1_ and P_3_, were utilized for different inlet and outlet configurations. The bottom channel outlet was kept closed throughout the experiments to ensure that water flow passed through the membrane. Pressure sensor P_1_ was positioned between the syringe pump and the microfluidic device, while P_3_ was placed after the microfluidic device. Water flowed from the syringe pump to the microfluidic device through P_1_, whereas P_3_, being open to the atmosphere, was located downstream. This process was repeated for PEGylated membranes, immediately following the completion of grafting when the system was still fully hydrated.

### Liposome & EV Staining

200 nm liposomes (Encapsula NanoSciences, Brentwood, TN, USA, CEP-500, Lot: CEP500-12082022) at a stock concentration of 1×10^13^ nanoparticles (np)/mL were serial-diluted using PBS to obtain a working concentration of 4×10^8^ np/mL. Utilizing nanotracking analysis (Malvern Nanosight NS300) we determined that the true mean size of the liposomes used were closer to 170 nm in size.

Similarly, HUVEC-derived extracellular vesicles from internal experiments were utilized. Nano-tracking analysis confirmed the average size of the EVs to be 145 nm and working solutions of 1×10^8^ np/mL were prepared.

A ratio of 1:100 Cell-Mask Orange, CMO, (Invitrogen, C10045, Lot#2738400) to nanoparticles (liposome or EV) in the working solution was used for labeling. Following sufficient mixing, the nanoparticles and CMO were incubated for an hour at room temperature prior to use in experiments.

## Funding

This work was supported by the National Institute of General Medical Sciences (NIGMS) through grants R21GM146156 and R35GM153461 to TRG. Additional support was provided by the Corporate Research department of Sartorius.

## Competing interest statement

MD is an employee of Sartorius Stedim North America, which provided financial support for this study. MD was involved in the design of the experiments, but not the data collection or analysis.

## Supporting Information

**Figure S1:**
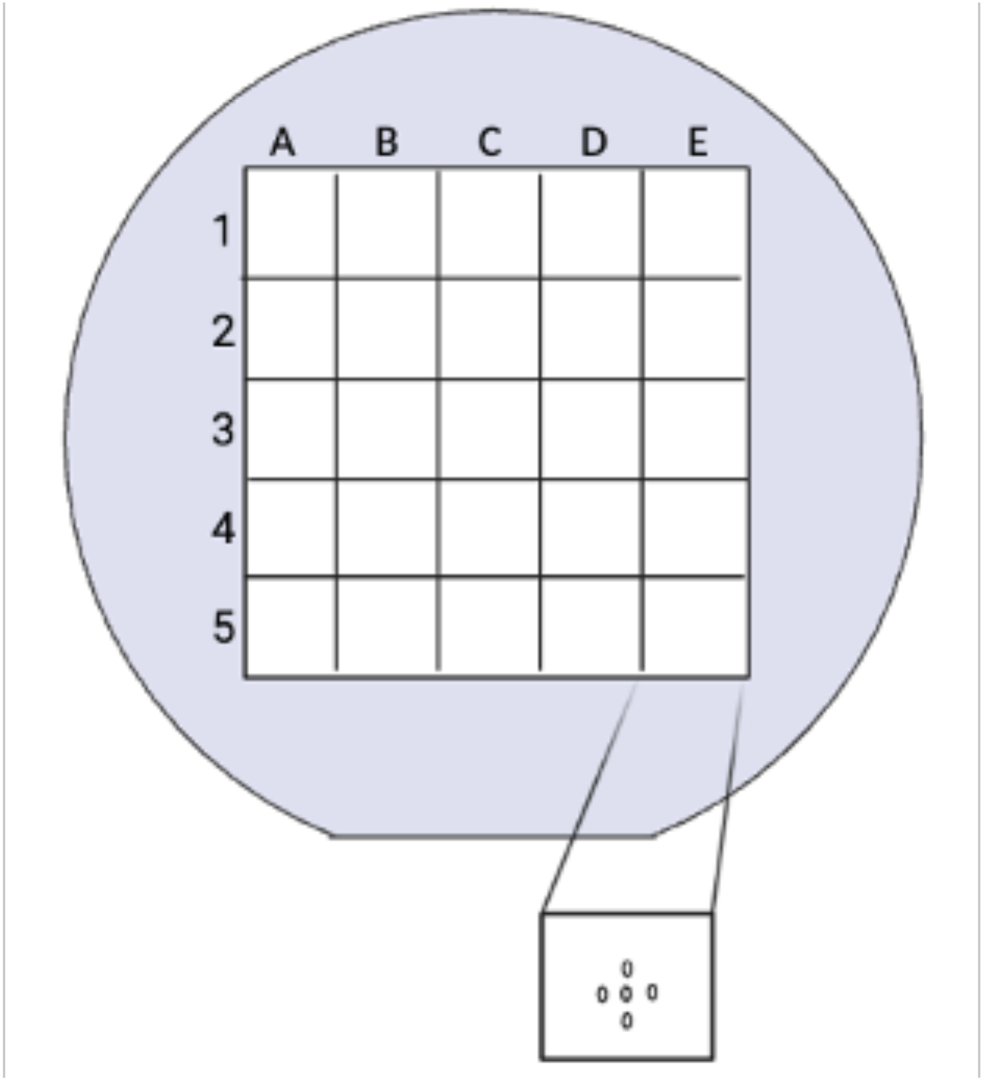
The indexed physical template, designed to mimic the wafers, for measuring the thickness of the parylene at a consistent locations across all wafers. The small square with five spots represents the number of measurements taken per square using the Nano-Spec. The indexed physical template was divided into a grid of 5×5 squares, labeled A to E along the x-axis and 1 to 5 along the y-axis. This structure was used to determine parylene thickness at specific locations. For each square, five measurements were taken using the NanoSpec, and the average was calculated. This average serves as the representative thickness for that spot. A fabricated shape to mimic the wafer shape was used to ensure that the NanoSpec measurements were taken at the same location on all wafers (Figure S2).

**Figure S2:**
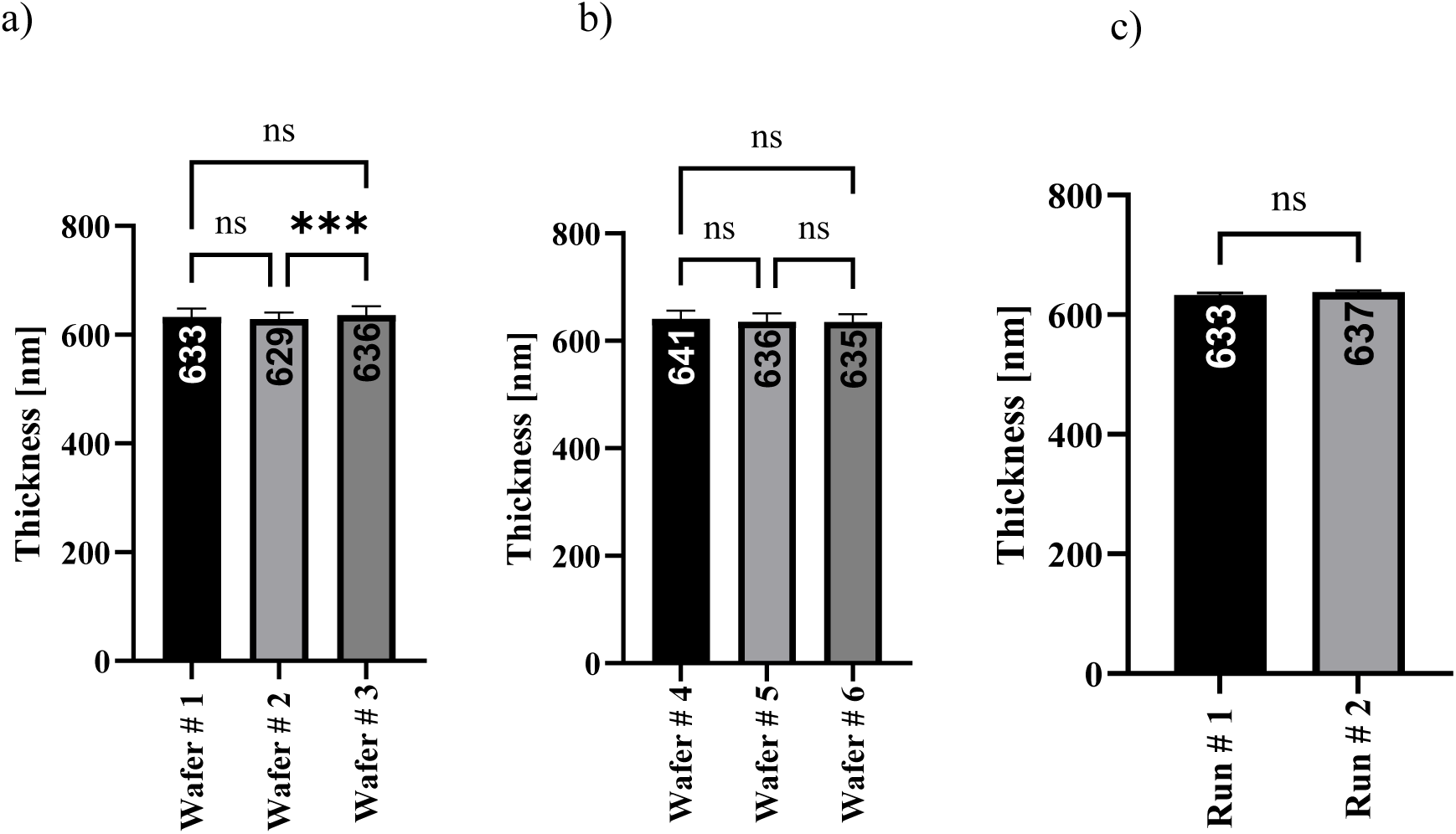
a and b) Wafer-to-wafer differences in parylene thickness within a single run; each run consisted of three wafers with 125 measurements per wafer (n = 125). c) Run-to-run differences in parylene thickness, comparing two independent runs, each comprising three wafers (n = 3 per run).

**Figure S3:**
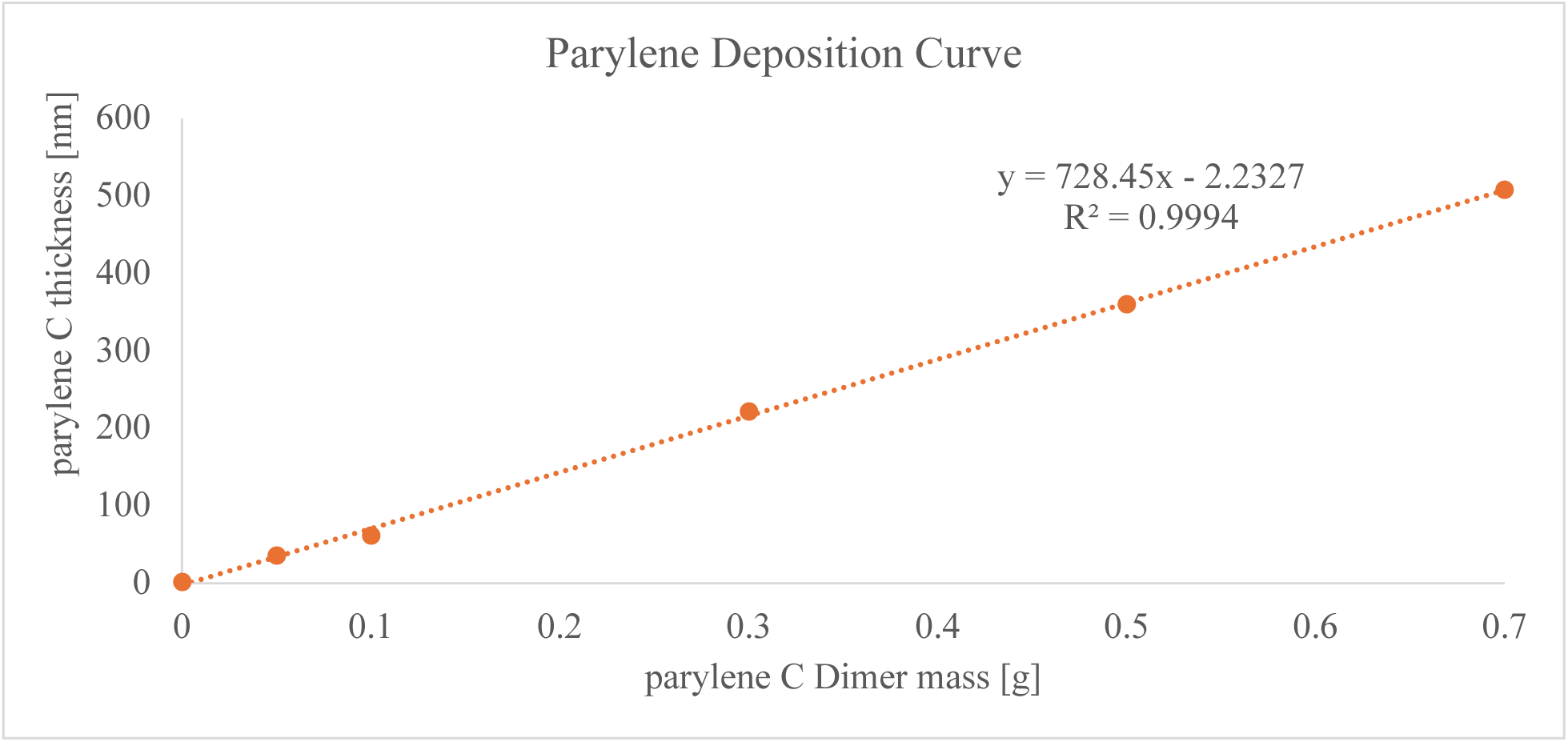
Effect of different parylene amounts on the film thickness and pore size. Deposition curves from various runs illustrate the relationship between the deposited parylene mass (x-axis) and the resulting film thickness (y-axis) and show how the deposited mass influenced the top pore size.

**Figure S4:**
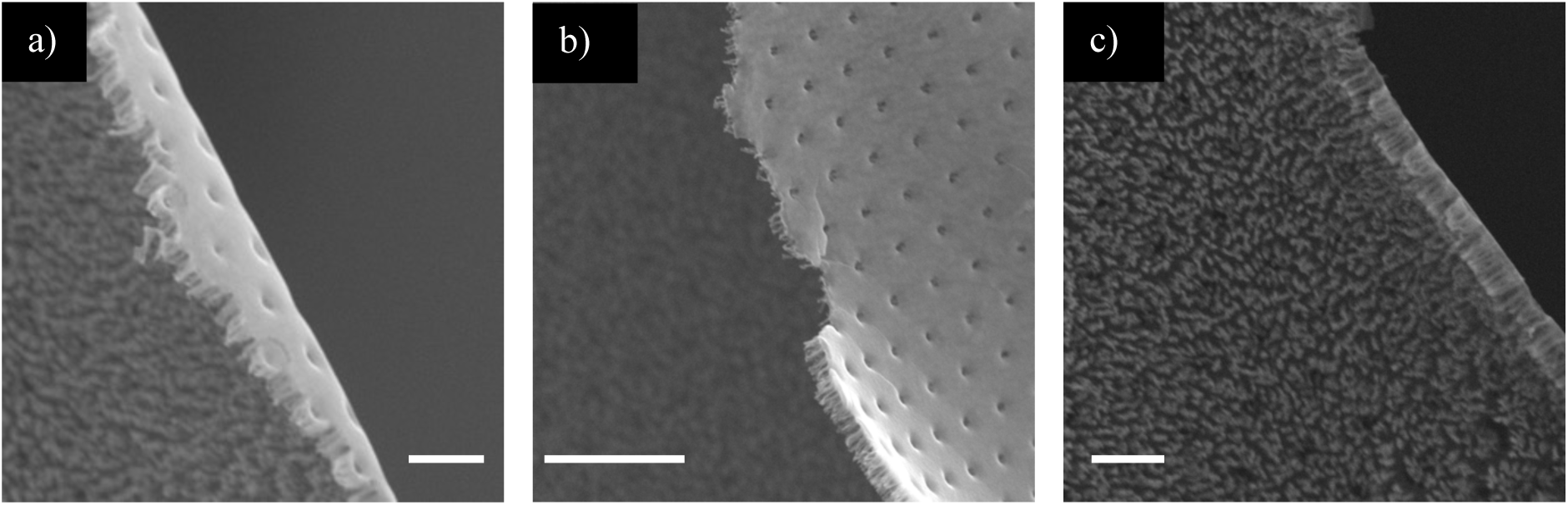
SEM images of parylene after etching, showing the nanoforest phenomenon. All images display the nanoforest structure from different viewing angles. a) Top, side, and bottom views; b) top and side views; c) bottom and side views. Scale bars = 2 µm.

**Figure S5:**
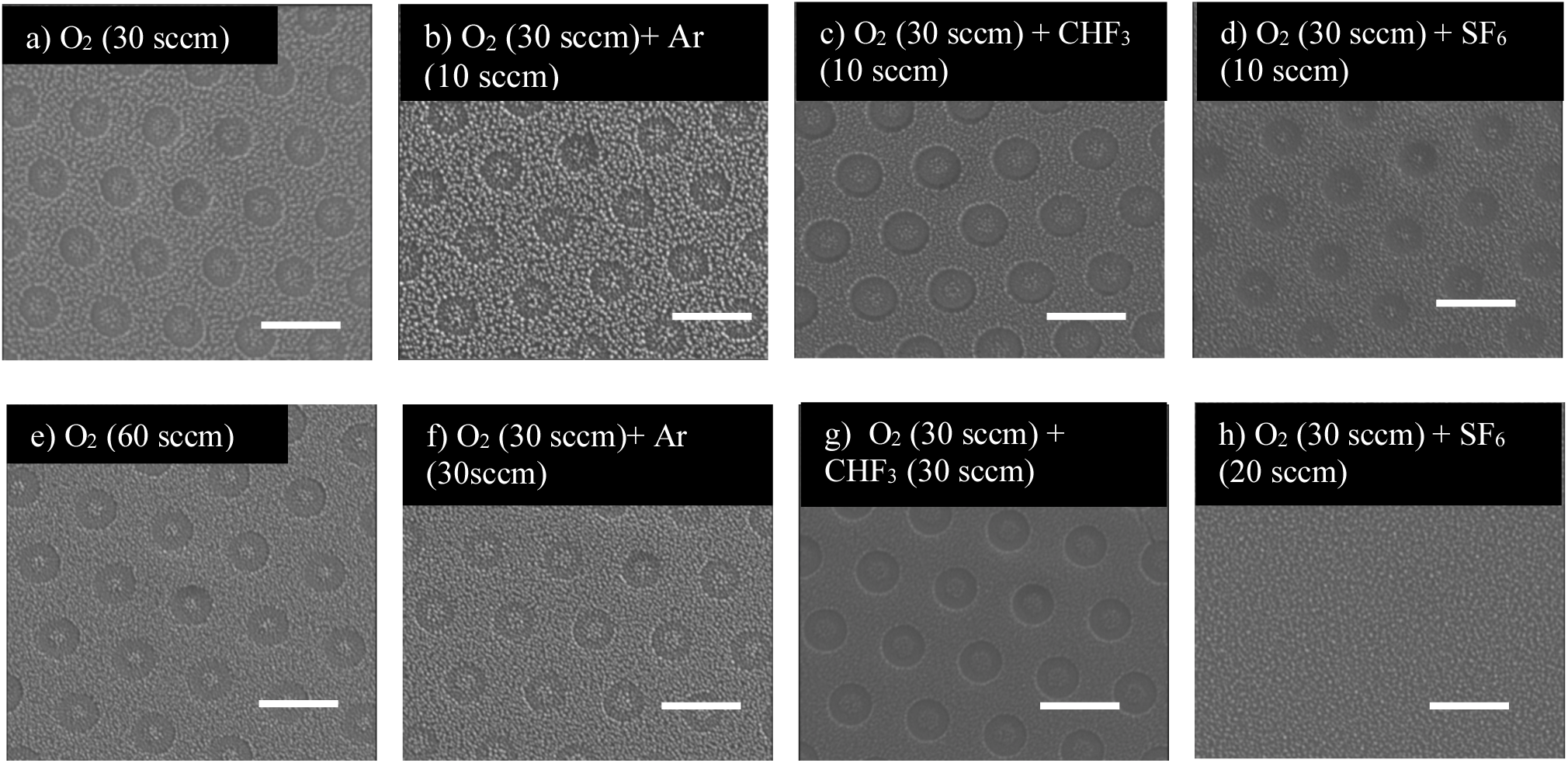
SEM images of parylene showing etch differences under various gas combinations and flow rates, aimed at reducing the nanoforest issue. **SEM images for a &** e) O_2_ (30 and 60 sccm), b & f) O_2_ (30 sccm)+ Ar (10 and 30sccm) c & g) O_2_ (30 sccm) + CHF_3_ (10 and 30 sccm) d & h) O_2_ (30 sccm) + SF_6_ (10 and 20 sccm). Scale bars = 2 µm During parylene etching with O_2_, we observed the formation of nanoforest structures. Figures S5a and S5e show the effect of O₂ flow rates at 30 and 60 sccm, respectively. Figure S5e illustrates that the nanoforest phenomenon increases with higher O_2_ flow rates, which led us to choose a lower O_2_ flow rate. To further reduce the nanoforest formation, we conducted a series of rigorous experiments. We introduced various gases, including argon (Ar), CHF_3_, and SF_6_, alongside low-flow O_2_ at flow rates of 10 and 30 sccm. Our approach revealed that CHF_3_ and SF_6_ outperformed the other gases at higher flow rates. However, we noticed that the etch rate became unstable at these high flow rates for both CHF_3_ and SF_6_. We determined that a stable flow rate of approximately 8 sccm was optimal for both CHF_3_ and SF_6_. For subsequent experiments, we selescted flow rates of 30 sccm for O_2_ and 8 sccm for CHF_3_, based on their effectiveness in reducing the nanoforest phenomenon while maintaining etching stability.

**Figure S6:**
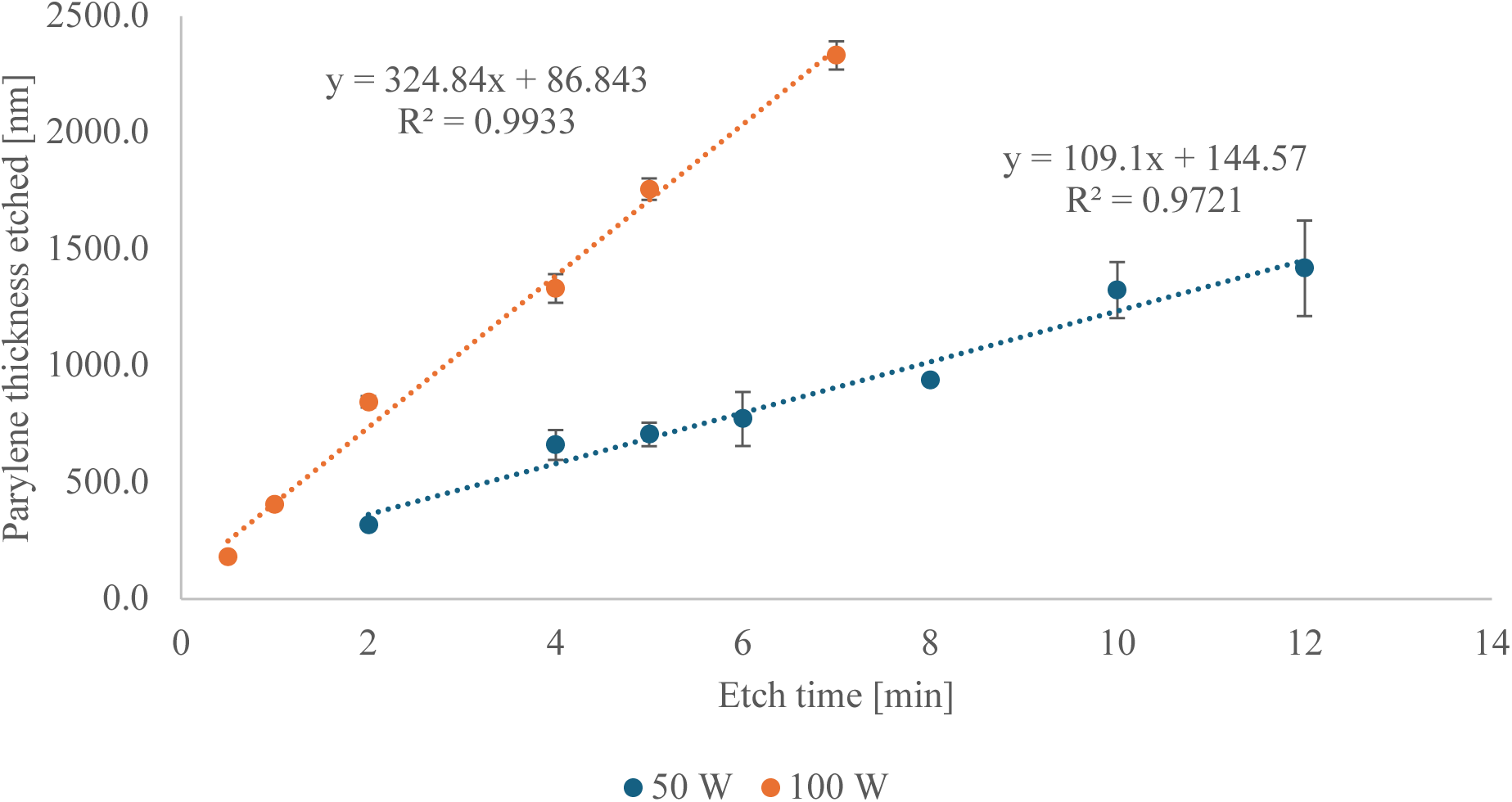
Effect of etching power (50 and 100 W) on the etch rate. Etch rates for two different power levels 50 W (blue) and 100 W (orange), showing how increased power accelerated the etching process.

**Figure S7.**
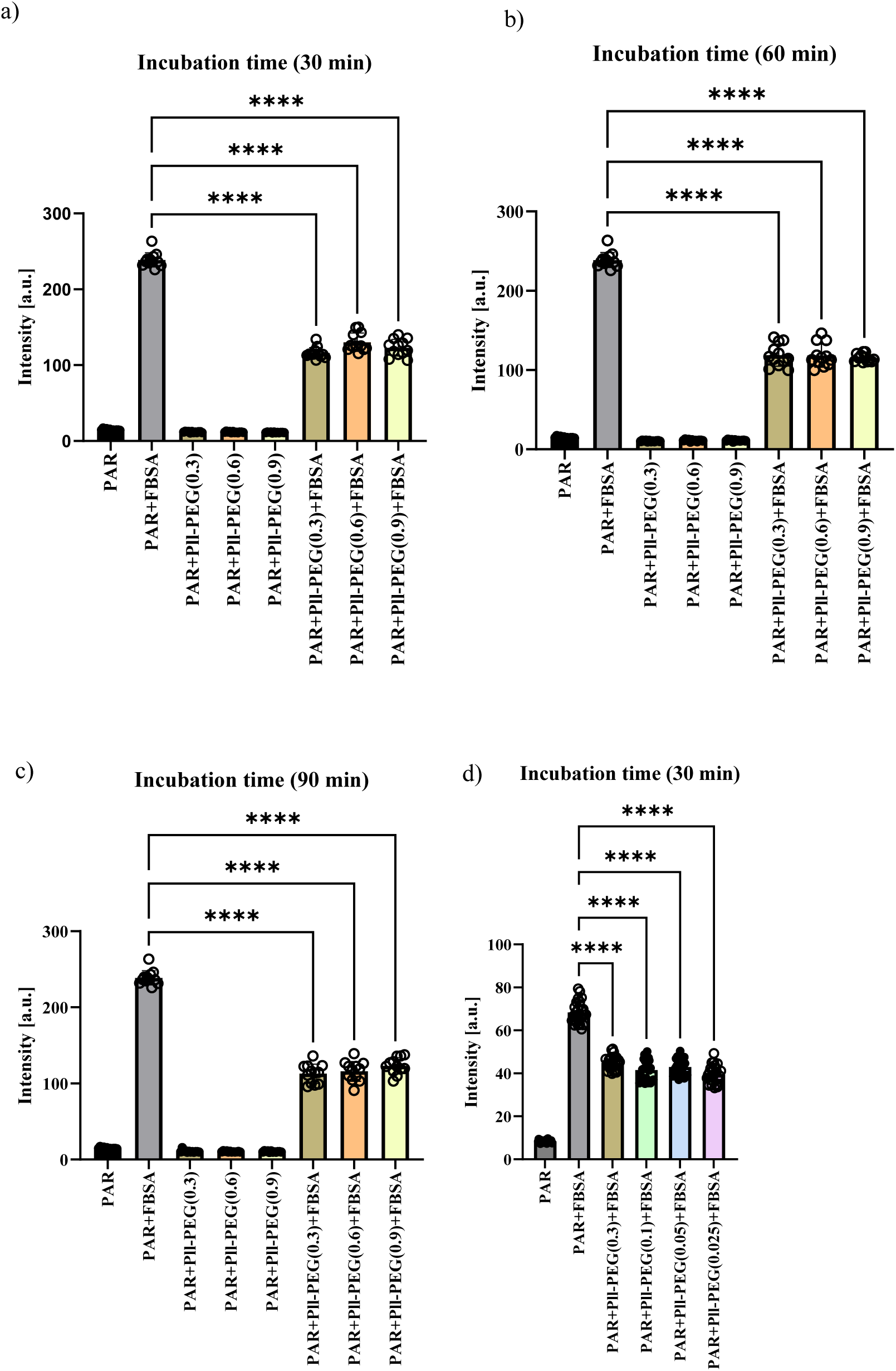
Optimization of PLL-PEG concentration and incubation time. a) PLL-PEG at 30 min incubation time for 0.3, 0.6 and 0.9 mg/mL. b) PLL-PEG at 60 min incubation time for 0.3, 0.6 and 0.9 mg/mL. c) PLL-PEG at 90 min incubation time for 0.3, 0.6 and 0.9 mg/mL. d) Lower PLL-PEG concentrations were then tested using a 30-minute incubation, **** = p < 0.0001. Initial experiments assessed PLL-PEG concentrations of 0.3, 0.6, and 0.9 mg/mL with incubation times of 30, 60, and 90 minutes to evaluate protein fouling (Figure S7 a-c). All tested conditions significantly reduced f-BSA adhesion, with no notable differences across concentrations or incubation durations. To improve grafting efficiency, we subsequently investigated lower PLL-PEG concentrations with a fixed 30-minute incubation (Figure S7d). This optimization identified 0.025 mg/mL for 30 minutes as the most time- and resource-efficient condition.

**Table S1:** Comparison of Brush Lengths on Parylene Under Different Concentrations and Hydration States. PLL-PEG at concentrations of 0.025 mg/mL and 0.9 mg/mL exhibited comparable brush lengths on flat parylene substrates under dehydrated conditions, with average lengths of 33.16 ± 5.18 nm and 34.81 ± 3.21 nm, respectively, suggesting that concentrations within this range are effective. The relatively large standard deviation is hypothesized to arise primarily from the intrinsic nanotexture and surface roughness of parylene C ^20^. Under hydrated conditions, brush lengths remained consistent with those measured in the dehydrated state, yielding values of 31.57 ± 4.52 nm and 35.79 ± 3.06 nm for the 0.025 mg/mL and 0.9 mg/mL concentrations, respectively

| PLL-PEG Concentrations [mg/mL] | Dehydrated | Hydrated |
| --- | --- | --- |
| **0.025** | $33.16 \pm 5.18$ nm | $31.57 \pm 4.52$ nm |
| **0.900** | $34.81 \pm 3.21$ nm | $35.79 \pm 3.06$ nm |

**Figure S8:**
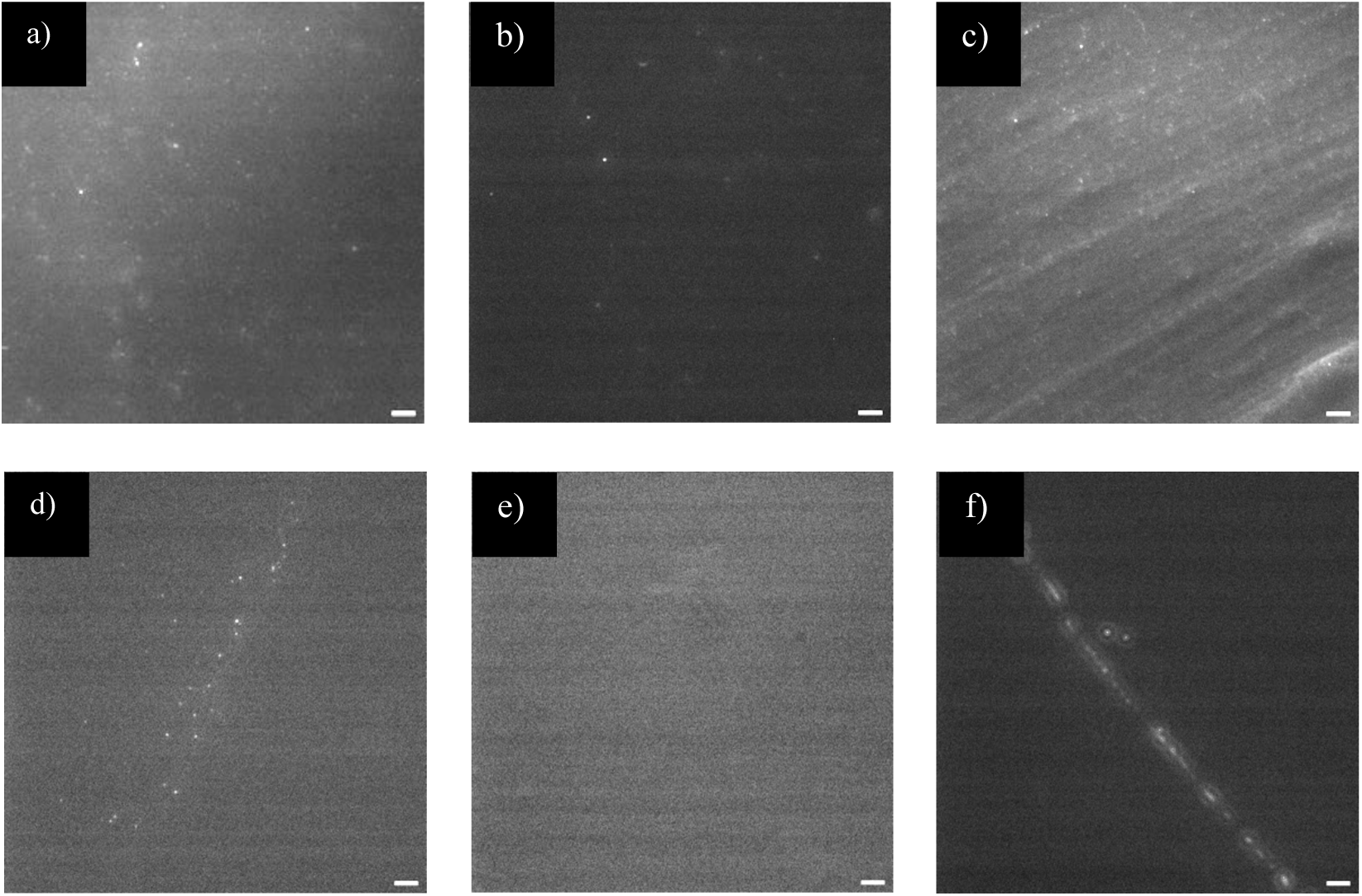
0.05 and 0.1 µm beads on glass slide. a-c) 0.05 µm beads on a glass slide collected from outlet 3 after passing through PLL-PEG–coated or uncoated membranes, a) 0.05 µm beads passed through the uncoated membrane, b) 0.05 µm beads passed through the PLL-PEG–coated membrane, c) 0.05 m beads control. d-f) 0.1 µm beads on a glass slide collected from outlet 3 after passing through PLL-PEG–coated or uncoated membranes, d) 0.1 µm beads passed through the uncoated membrane, e) 0.1 µm beads did not pass through the PLL-PEG– coated membrane, f) Control: 0.1 µm beads on glass slide. Scale bars= 10 µm.

**Figure S9:**
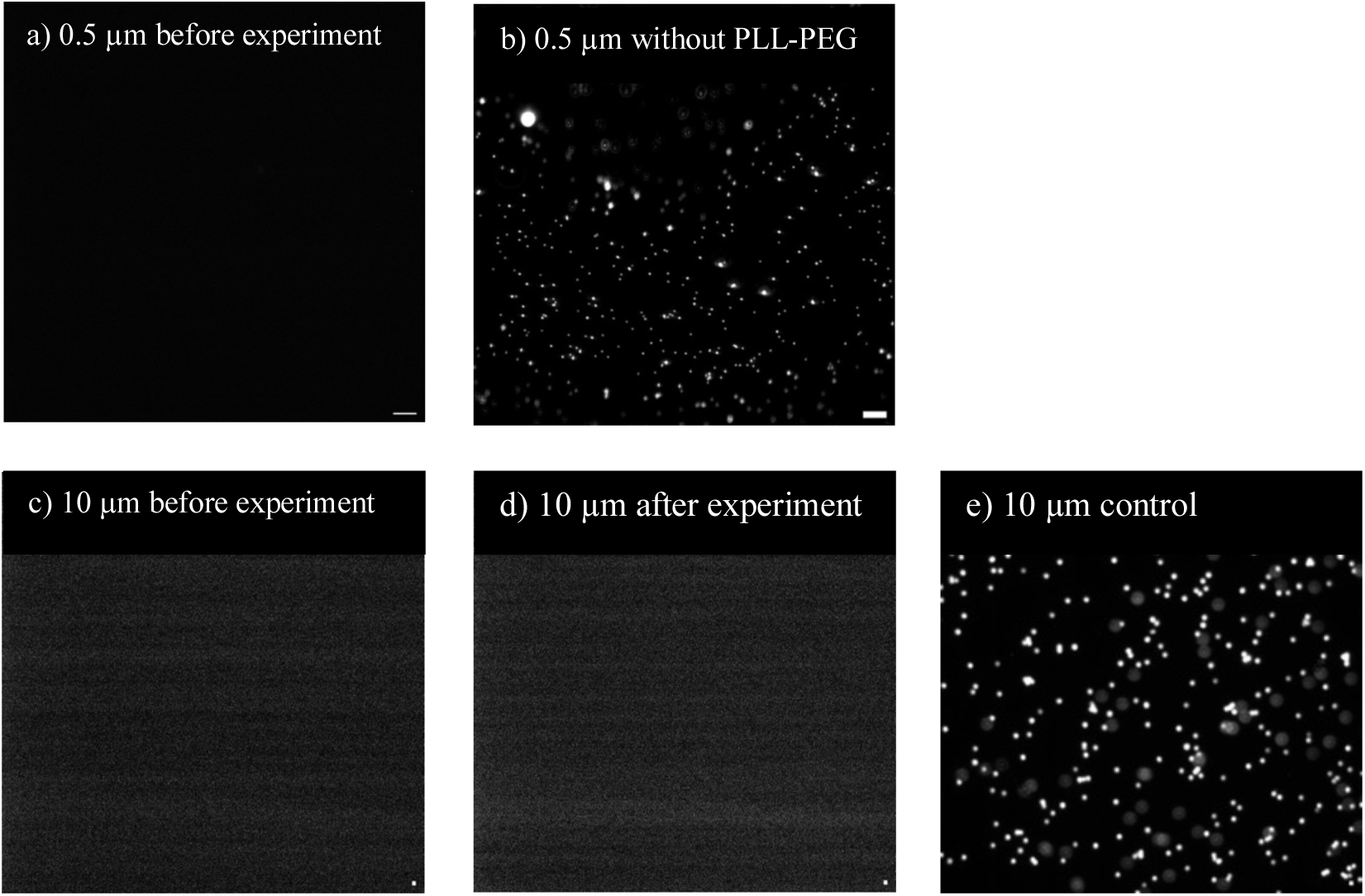
Fluorescence imaging of 0.5 µm and 10 µm beads on the membrane. a, b) 0.5 µm beads on the membrane without PLL-PEG coating, a) Membrane before beads introduction using pump, b) 0.5 µm beads captured on the membrane after the capturing step. c-e) 10 µm beads on the membrane without PLL-PEG coating, c) the membrane before beads introduction using pump, d) no 10 µm beads observed on the membrane after the capturing step, e) Control: 10 µm beads sandwiched between cover glass and glass slide. Scale bars= 10 µm.

**Figure S10:**
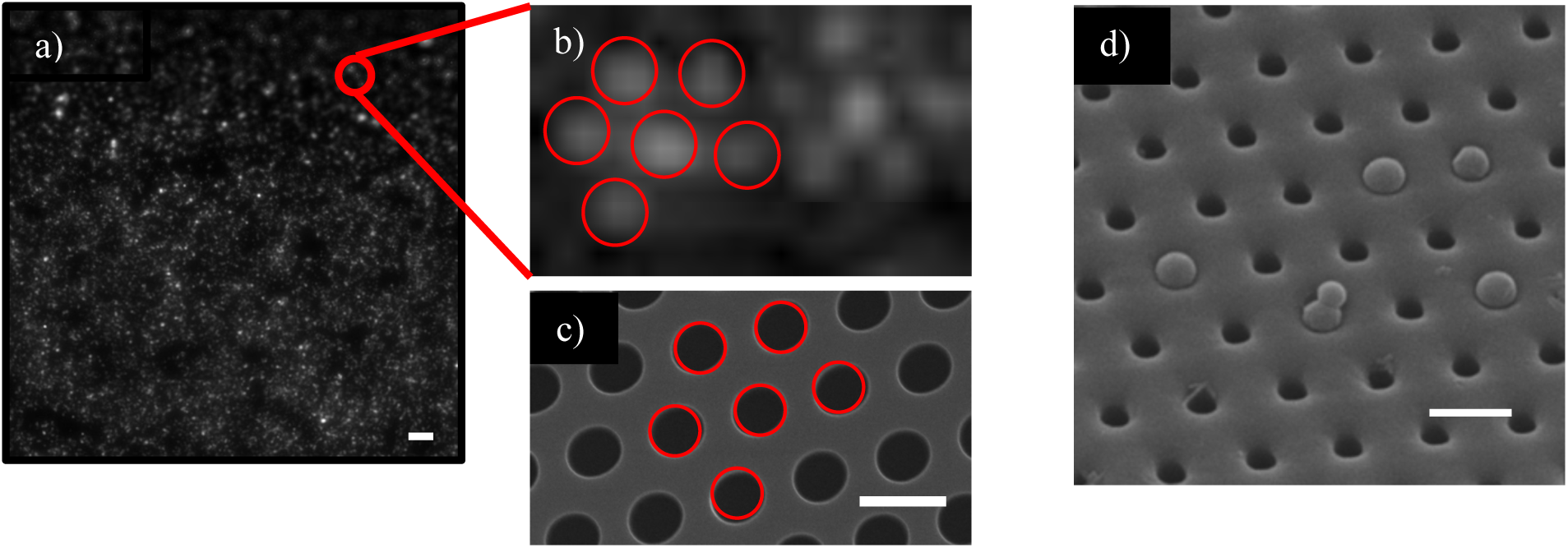
Beads orientation on the membrane. a) 0.5 µm beads were captured on the membrane, as determined through fluorescence microscopy, scale bar =10 µm. b) The orientation of fluorescent beads captured on the membrane, with red circles indicating their positions. c) SEM imaging of the nanopocket membrane mask suggests beads are successfully and specifically captured in pores, when comparing bead distribution in (b) to the red circles in (c), scale bar =2 µm. d) SEM imaging further proved that the 0.5 µm beads are captured by the nanopocket membrane, scale bar =1 µm

**Figure S11:**
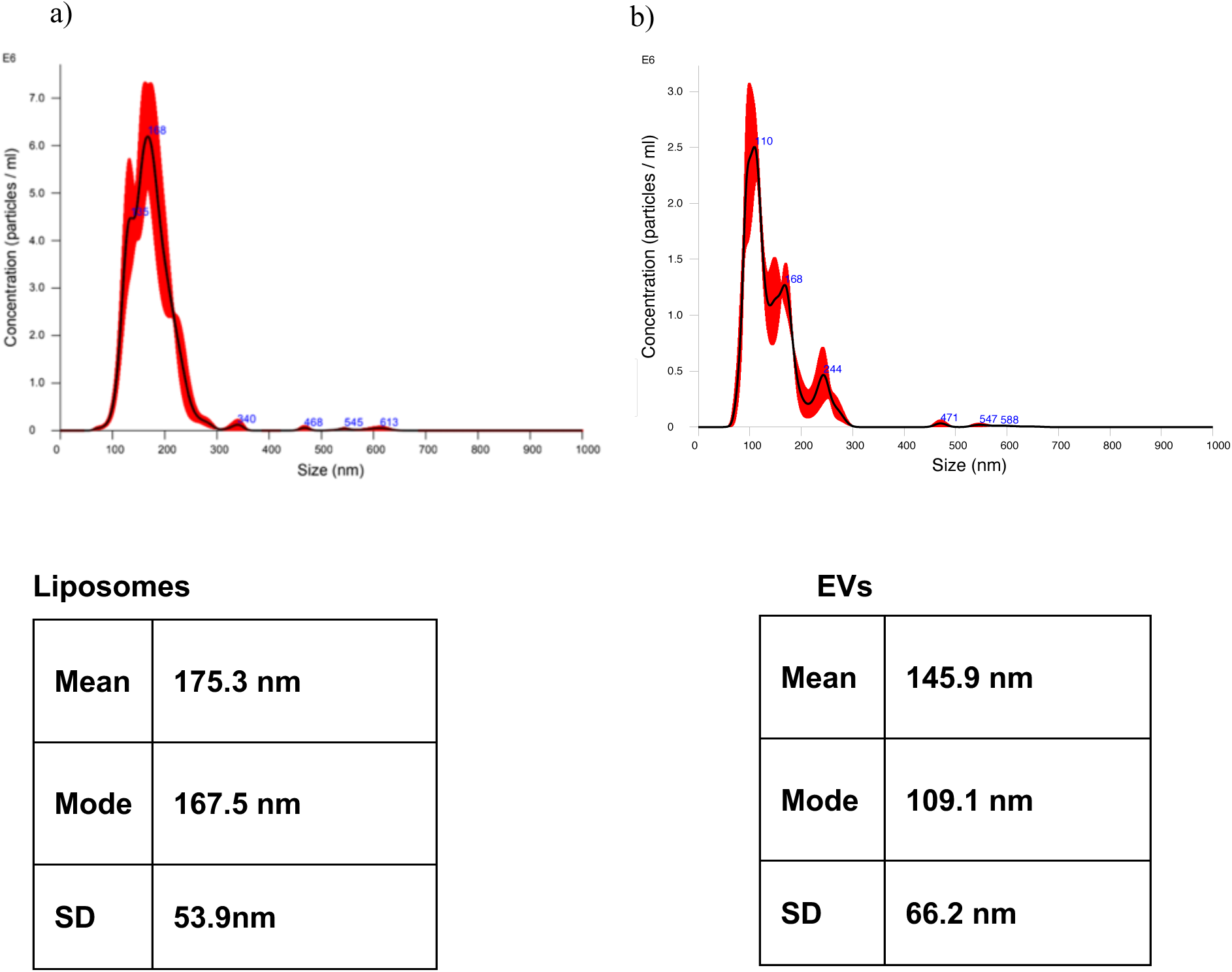
Nanoparticle Tracking Analysis (NTA) of liposomes and extracellular vesicles (EVs). a) Liposomes exhibited a mean diameter of approximately 175 nm. b) EVs exhibited a mean diameter of approximately 146 nm.

**Figure S12.**
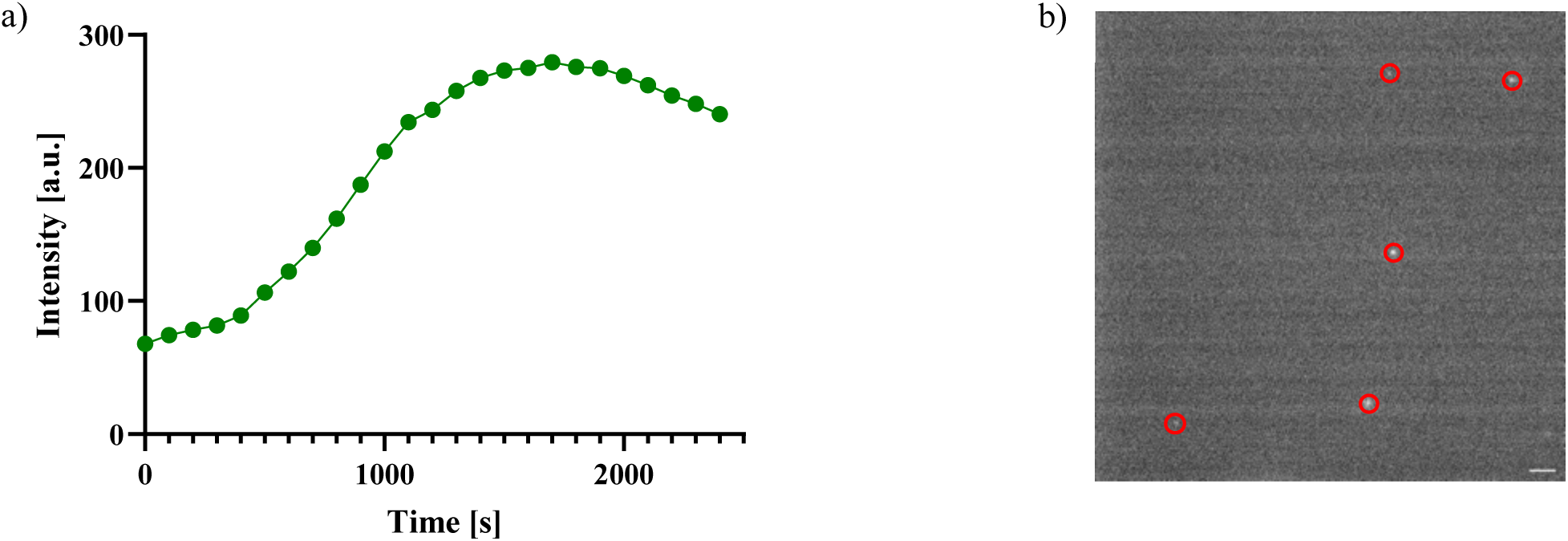
f-BSA permeability and EV fluorescence imaging on glass. a) f-BSA intensity increased over time in channel 3, indicating that the protein passed through the coated membrane to reach the channel. b) Fluorescently labeled EVs appeared immobilized between the cover glass and the slide, circles indicated EVs, scale bar= 10 µm

